# The Yeast Lifespan Machine: a microfluidic platform for automated replicative lifespan measurements

**DOI:** 10.1101/2022.02.14.480146

**Authors:** Nathaniel H. Thayer, Michael Robles, Jun Xu, Elizabeth L. Schinski, Manuel Hotz, Robert Keyser, Alfred Millett-Sikking, Voytek Okreglak, Jason V. Rogers, Adam J. Waite, Bernd J. Wranik, Andrew G. York, R. Scott McIsaac, Daniel E. Gottschling

## Abstract

The budding yeast, *Saccharomyces cerevisiae*, has emerged as a model system for studying the aging processes in eukaryotic cells. However, the full complement of tools available in this organism has not been fully applied, in part because of limitations in throughput that restrict the ability to carry out detailed analyses. Recent advances in microfluidics have provided direct longitudinal observation of entire yeast lifespans, but have not yet achieved the normal scale of operation possible in this model system. Here we present a microfluidic platform, called the Yeast Lifespan Machine, where we combine improvements in microfluidics, image acquisition, and image analysis tools to increase robustness and throughput of lifespan measurements in aging yeast cells. We demonstrate the platform’s ability to measure the lifespan of large populations of cells and distinguish long- and short-lived mutants, all with minimal involvement of the experimenter. We also show that environmental pH is capable of significantly modulating lifespan depending on the growth media, highlighting how microfluidic technologies reveal determinants of lifespan that are otherwise difficult to ascertain.

## Introduction

The budding yeast, *Saccharomyces cerevisiae*, is a model system for developing fundamental understanding of eukaryotic cell biology, including cellular aging. These cells divide asymmetrically, where during each division a mother cell “buds” to produce a daughter cell. The number of daughter cells that each mother cell can produce is limited and defines the cell’s Replicative Lifespan (RLS) (1). As a mother cell produces more and more daughter cells, cellular processes and functions begin to deteriorate, which ultimately results in the cell’s death (reviewed by Denoth et al. (2)). Comparative studies of aging processes across model organisms have identified shared molecular pathways that decline during aging, providing evidence that aspects of the mechanisms of aging are conserved across eukaryotes(3).

Aging is recognized to be the complex interaction between genetic variation and environment. Genetic and environmental perturbations have been identified that modulate the RLS of yeast cells. However, a full exploration of the interplay between growth environment and the importance cellular pathways have on RLS has been hampered by technical challenges. Specifically, the traditional RLS assay requires manual separation of daughter cells from the mother cell after every division, an incredibly tedious process (4). Despite these challenges, McCormick et al., performed a cursory RLS screen of the entire Yeast Non-essential Gene Deletion Collection by micromanipulation (5, 6). This study revealed many important genetic modulators of RLS, but due to the thousands of hours needed to obtain sufficient measurements, this screen suffered from a lack of discriminatory power to identify a comprehensive list of genes impacting aging. Furthermore, the effort involved in this approach makes it unlikely that it can be practically used to explore additional genetic backgrounds or environmental conditions.

Several technologies have been developed to overcome the challenges of studying yeast aging by micromanipulation. Some approaches allow purification of aged cells from a large population and have enabled biochemical and cell-biological experiments in aging yeast cells (7–10). However, direct and longitudinal observation of phenotypes in a significant number of cells remained impossible until microfluidic designs were introduced to automatically separate daughter cells from mother cells (11, 12). This technique alleviates the need for constant human intervention and permits the use of high-resolution microscopy techniques to monitor the aging process. Despite many research groups developing different devices and employing several distinct strategies, no single device has been universally adopted by the field (comprehensively reviewed by Jo et al. and Chen et al.; experimentally explored by Gao et al. (13–15)).

An emerging challenge of using microfluidic devices coupled with high-resolution microscopy techniques is the development of automated techniques to determine the RLS from the large volume of raw data. While some devices employ chemical/genetic methods to determine lifespan by counting the number of arrested daughter cells at the end of the experiment (16), most systems require constant longitudinal imaging to generate time-lapse movies of the entire lifespan of each cell. Traditional image analysis techniques are inadequate to automate RLS quantification from these movies due to the heterogeneity of budding patterns, cellular morphologies and temporary cell crowding observed in microfluidic devices. Incorporating fluorescent markers to facilitate mother-cell tracking or to mark the completion of each cell division has been used to facilitate human annotation or even automate some analyses (17, 18). However, these strategies are difficult to scale because of the strain engineering, sample preparation, or additional imaging modalities needed. Another significant hurdle is the annotation and analysis of the longitudinal lifespan movies. Simply, these devices already produce more data than any researcher could hope to look at, let alone annotate. Fortunately, this problem of having too much data to effectively analyze by traditional methods is well suited for automated approaches using computer-vision/machine-learning. Here, we describe the development of a microfluidics platform, called the Yeast Lifespan Machine, that overcomes the challenges of previously described methods to accurately quantify RLS in yeast. The Yeast Lifespan Machine leverages microfluidics to robustly collect lifespan data of high temporal resolution from large populations of aging cells (approximately 800 per channel) in a system with 24 independent channels, which can contain distinct genetic or environmental conditions. We developed a novel computer-vision strategy for extracting RLS data from longitudinal images of aging cells in the Yeast Lifespan Machine. We demonstrate our system’s ability to recapitulate the effects of known long- and short-lived mutants, and identify an environmental condition capable of modulating lifespan.

## Results

### Yeast Lifespan Machine Microfluidic Device

The Yeast Lifespan Machine is based upon a multi-layer PDMS microfluidic device with 24 discrete channels. Each channel contains an array of PDMS features specifically designed to retain aging mother cells while daughter cells are washed away (Fig.1A). Consequently, it is capable of measuring the replicative lifespan for a particular genotype in a specific environmental condition. Each channel has three access ports – one used as media input, one for cell input, and one for output of waste and daughter cells (Fig.1B). While other devices have been able to simplify experimental setup by sharing one or more of these ports between multiple microfluidic channels, the Yeast Lifespan Machine provides greater overall experimental robustness (19). By keeping each microfluidic channel separate, any one failure (a clog, contamination, etc.) is isolated, and thus does not compromise data collected from the other, separate, microfluidic channels.

**Fig. 1.**
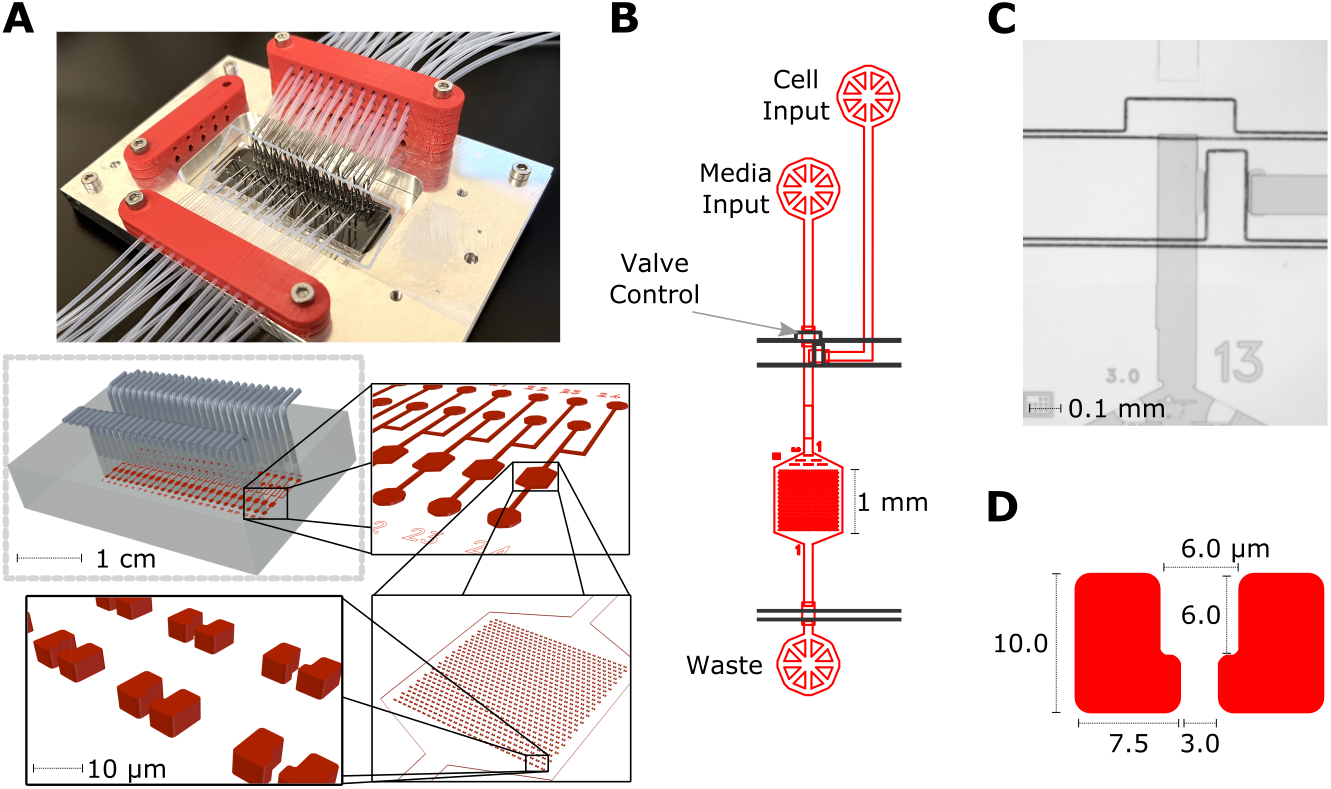
Yeast Lifespan Machine microfluidic device. A) The Yeast Lifespan Machine microfluidic device is a PDMS device with 24 discrete channels, each with an array of 850 cell “catchers”. B) Each channel has 3 ports. One used to input cell cultures. One used to provide fresh media for the duration of the experiment. One used to remove waste from the microfluidic channel. These 3 ports are connected by the fluidic channels (red). Flow in these fluidic channels are controlled by a series of microfluidic valves (black boxes), actuated by overlapping control channels in the valve layer (black). C) Microfluidic valves are actuated by pressurizing the control channel above the fluidics channel, completely collapsing and sealing the channel below. In this example image, both valves are closed, blocking media flow in the channel. D) Dimensions of catcher features.

Precise control of fluid flow throughout the device is controlled by a series of microfluidic valves, operated by pressurizing a separate layer of the device, reversibly collapsing the microfluidic channel below (Fig.1B,C) (20). These valves allow for a relatively simple layout of microfluidic flow channels, yet completely isolate media input ports from cell loading ports. This configuration greatly reduces the risk of cells accumulating in any part of the microfluidic device, which could negatively impact the experiment. While other devices have achieved suitable control with creatively designed channel layouts, we have found that the increased robustness provided by the sealed valves is worth the additional steps of design and fabrication (12, 19).

Each channel has a large array of approximately 850 PDMS features designed to retain aging yeast cells. These cell “catchers” or “traps” were inspired by a previously described device (the HYAA chip) and are designed to allow budding of the mother cell in multiple directions without the cell being lost from the trap (“washing out”), while daughter cells are removed by a constant media flow (Fig.1D)(19). While original cells are preferentially retained by these catchers and often retained until death, some cells escape or are crowded by other cells at various times during the experiment and must be censored (Fig.S1). Within each channel, cell catchers are in a 29×30 array, which measures approximately 1mm x 1mm, and can be tiled with 9 fields-of-view on our microscope systems.

### Image Acquisition, Processing and Computer Vision

The overall design of our image analysis pipeline is illustrated in Figure 2 and specific aspects of this pipeline are described in the following sections.

**Fig. 2.**
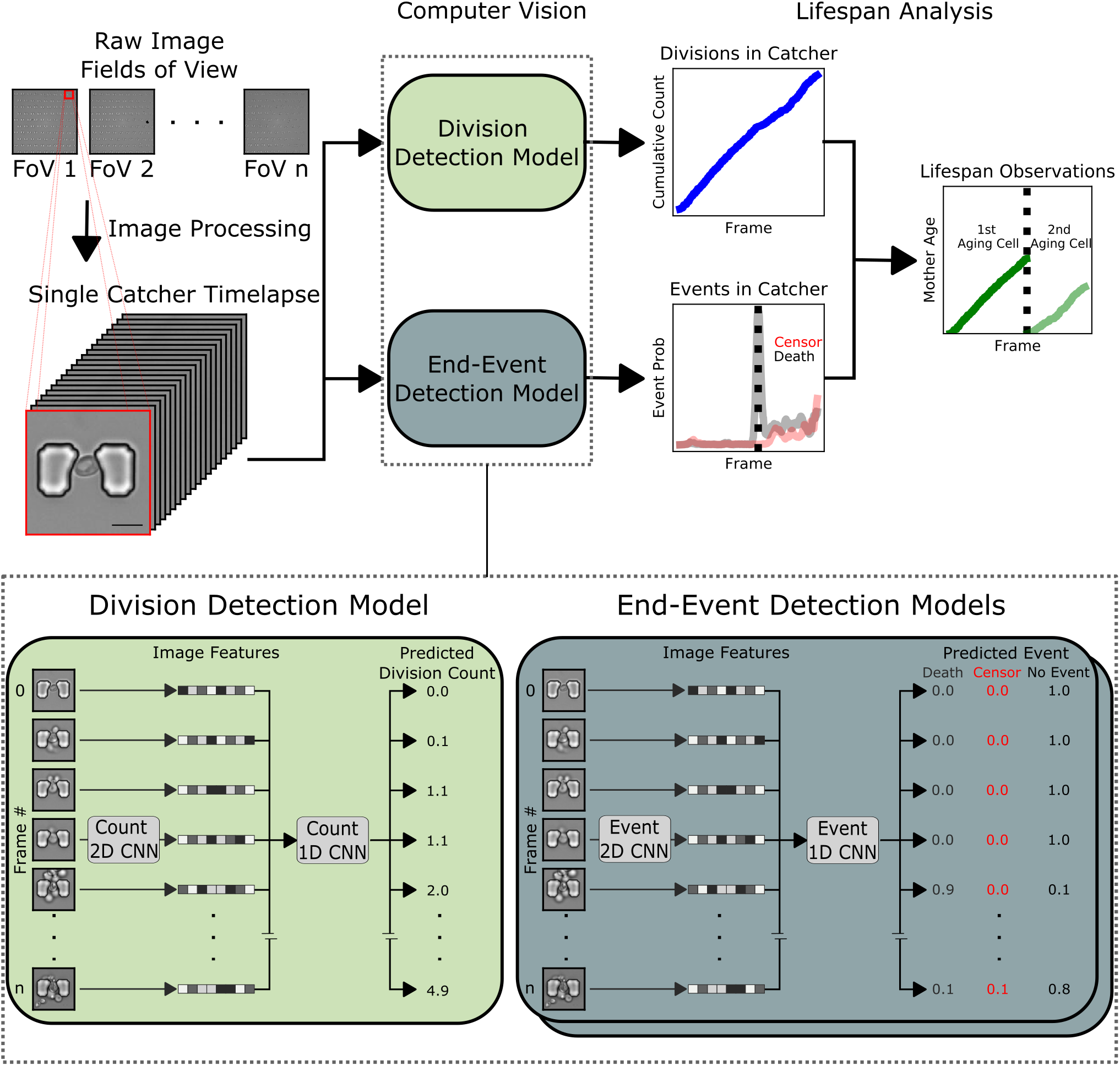
Image processing and computer vision pipeline. The raw images of each field-of-view are corrected for drift, then processed into smaller image patches, each containing a single catcher. The time series of each catcher are then analyzed by two computer vision models. One model detects the number of cell divisions that occur in the time series and one model determines when aging mother cells have died or escaped the catcher (censored). The output of each model is combined into lifespan observations. Insets: The architecture of each model can both be summarized by a 2D image feature extraction layer and a 1D dilated convolutional layer, capable of using information from surrounding frames to make predictions on i) division events or ii) end-events (i.e. mother cell escape from, or death within the catcher).

#### Image Acquisition

Time-lapse images are acquired using several instances of a custom-assembled, automated microscope (Fig.S2). Acquisition is controlled by a custom python application, allowing synchronization with microfluidic and environmental control hardware. These custom applications simplify experimental setup, handling of image data, collection of metadata, acquisition monitoring, and a number of other tasks that would have been difficult to automate with existing microscope control software (see materials and methods, supplemental material).

#### Image Processing

The raw images/data of each experiment can be considered as a six-dimensional array - in ‘PTZCYX’ order. ‘P’ for position or field-of-view, ‘T’ for time, ‘Z’ for z-slice, ‘C’ for imaging channel, and the ‘Y’ and ‘X’ dimensions of each image. A typical experiment to measure replicative yeast lifespan contains:

- 216 positions – 24 microfluidic channels, 9 fields-of-view per channel.
- 396 time points – taken at 15 minute intervals for 96 hours
- 5 z-slices – spanning 4μm (1μm spacing)
- 1 imaging channel – brightfield or DIC image
- 2048 Y and 2048 X pixels – the standard dimension of our sCMOS chip

Each field-of-view in a typical experiment represents about 16 GB of images (396T x 5Z x 1C x 2048Y x 2048X x 2 bytes/pixel = 16.6 gigabytes). Even when lossless compression is applied, the total data for all fields-of-view in a typical experiment can still total 0.5 - 1.5 TB of data. Given this large size, we process each field-of-view into smaller image patches that each contain a single cell-catcher. Each field-of-view is first processed by registering the ‘ZCYX’ data along the ‘T’ dimension, correcting for any thermal drift or imprecise stage movements during the experiment. A single ‘ZC’slice (shape: [396, 1, 1, 2048, 2048]) is used as a reference, and these corrections are applied to all other channels and z-slices. Next, the coordinates of each cell catcher are found by cross-correlating a template image of a catcher against the full image. Then a 256×256 window is selected around each cell-catcher. This process creates a 5D stack, typically with the dimensions [396, 5, 1, 256, 256], in ‘TZCYX’ order. While the resulting images are a more manageable 260 MB each, we often reduce the depth of the images to 8bit to cut the file size in half. Additionally, we compress these images in a tiled-jpg format that reduces the file size by 95-99%. While there is considerable loss of information at this compression step, the resulting images still preserve the information required for human or computer vision extraction of relevant lifespan information.

#### Computer Vision

To determine the replicative lifespan of each cell, three pieces of information are needed: i) which frames define the start and end of the cell’s lifespan, ii) whether the observation ended with a cell dying or washing out of the field-of-view, and iii) how many budding events occurred during the time the cell was observed.

Collecting data from a single microfluidic device can produce approximately 20,000 time-lapse images of individual lifespans. Even if an annotation task was relatively simple and took a human 15 seconds to annotate each image, it would still take nearly 87 person-hours of labor to annotate all of the images from the 24 channels in a single experiment. To take full advantage of all the data this system produces, we developed a high-throughput automated image analysis pipeline. We opted to train two separate models, the results of which can be combined to determine the lifespan of any given cell (Fig.2 inset). This allowed us to frame each as a relatively easy-to-annotate task and clearly define the problem that each model was trying to solve.

#### Division Detection Model

The first model was trained to predict the cumulative number of buds that have been produced by aging mother cells during the course of a timelapse movie. The input of this model was a 3D TXY stack of images, representing the full time-lapse movie of a single catcher. The first module of this model was a ResNet-like model to extract image features from each 2D slice of the timeseries (21). The output was flattened into 1D image features. Each set of 1D image features was then fed into the 1D convolutional neural network (CNN) Module, composed of a series of gated dilated convolution layers with residual connections (22), which is capable of using information from surrounding time points to determine if a division was likely to have occurred at each time point. The dilated convolution increases the receptive field size and allows a larger number of surrounding time points to be included.

This model was trained using approximately 10,000 annotated time-lapse series, optimizing for predicting the cumulative number of divisions that had occurred in each time series at each time point. To avoid the model learning the average budding rate of our cells and predicting bud counts by simply counting frames, we took two steps. First, we included training data from multiple growth conditions and genotypes that varied growth rate by 100-200%. Second, as a data augmentation step, we created artificial lifespan movies with random time points removed.

Overall, this model was able to predict the total number of divisions that had occurred in a movie with good accuracy (R^2^=0.89; Fig.3A). While not a perfect reproduction of the annotations, this error is similar to that we observed when comparing the annotations of two different annotators (R^2^=0.74; Fig.S3). In examining the difference between the predicted age and the annotated age at each frame of the movie (Fig.3B), the majority of events deviate from the annotated age by a small amount and are not biased to occur at any particular point in the time-lapse. There also does not seem to be a systematic over/under prediction of age by the model; the mean error is very near zero over all frames.

**Fig. 3.**
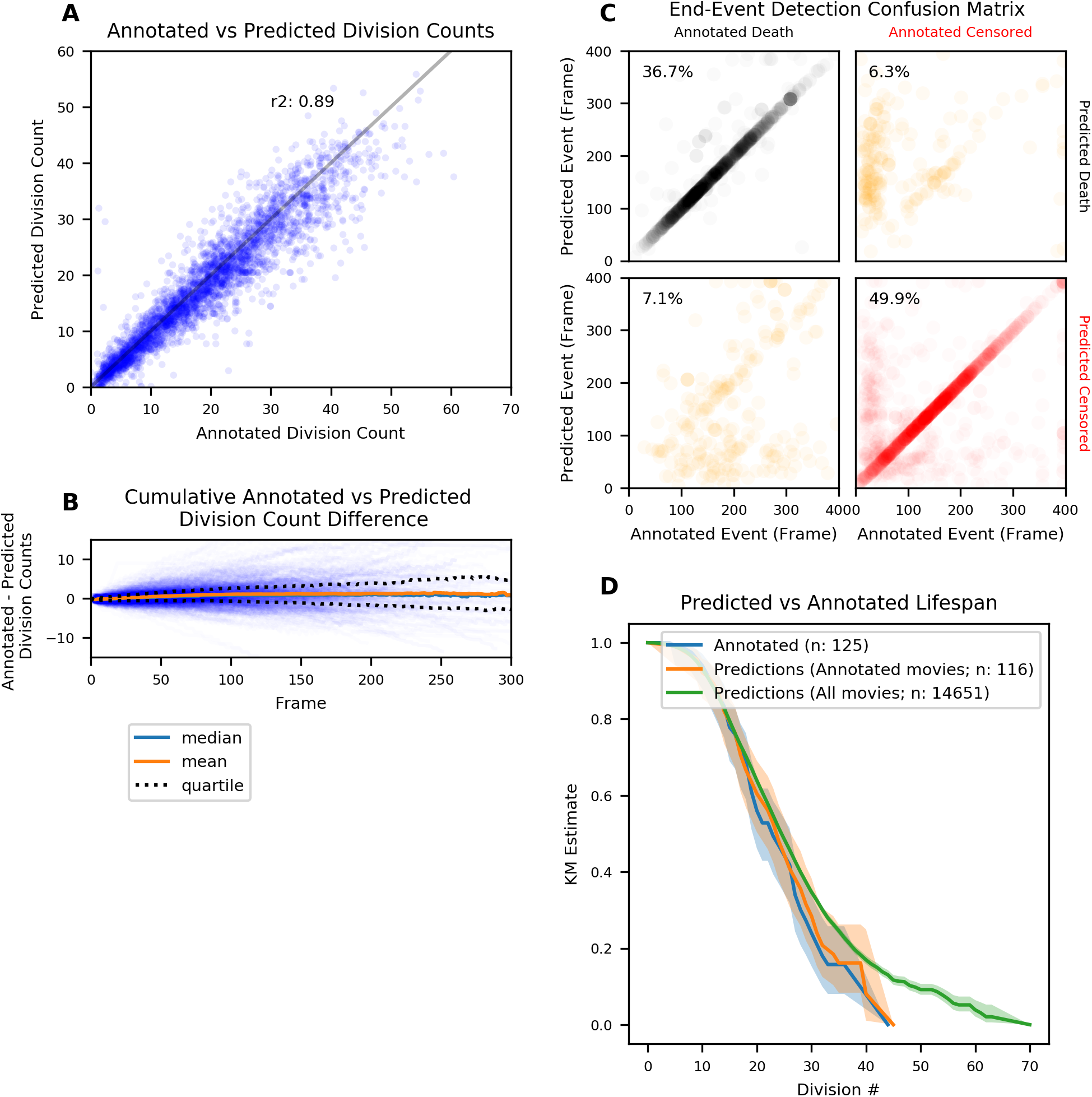
Computer vision validation. A) The performance of the model at predicting the final age of a cell, visualized as a scatter plot against ages measured by hand-annotation. B) The performance of the model throughout the lifespan of a cell, visualized as the difference between the predicted and annotated age. Each blue trace represents a unique lifespan. The median, mean and quartile ranges are also plotted. C) The performance of the event detection model visualized on a confusion-matrix / scatter plot. The placement of each point onto each quadrant of the confusion-matrix is determined by the annotated and predicted type of event. Each scatter plot indicates agreement on the frame at which the annotation or prediction is made. D) Kaplan-Meier survival estimates made from a set of time-lapse images that were both human annotated (blue) and predicted from computer vision pipelines (orange), or predictions made from all time-lapse images from the same experimental conditions (not just those that had been annotated).

#### End-Event Detection Model

The second model was trained to detect the end of each lifespan, replicating the annotations of when a cell died or was censored (e.g. cell washed out of the trap). The goal of the model was to recreate the specific type of annotated end-event near the frame in which it was annotated.

The model was an ensemble consisting of two models with very similar architectures, only slightly varying in their objective function. Again, the input of this model was a 3D TXY stack of images, representing the full time-lapse movie of a single catcher. The first module in each model was again an image feature extraction layer, this time utilizing a modified Unet architecture, which we found performs slightly better than a ResNet backbone (23). These image features were flattened, then fed into another 1D CNN module, similarly composed of a series of gated dilated convolution layers, to use information from surrounding time points to predict the likelihood of an event at each frame (22). The model was constructed to output a score for each frame related to the likelihood that an aging mother cell either died or censored at that frame. A threshold was established that if either score reached a value > 0.2, this event was predicted to have occurred, although the threshold is a tunable parameter. To account for the uncertainty of each annotation, we smoothed the probability of annotated frames to the surrounding frames using a gaussian kernel.

Overall, this model was able to accurately predict both the type and timing of the end-event for a majority of time-lapse movies of catchers. When comparing the annotated and predicted events in a confusion matrix (Fig.3C), the majority of events are plotted in either the upper-left or lower-right quadrant, indicating that the model accurately predicted the same type of event that was annotated. Furthermore, a majority of the events were predicted to occur on, or near, the same frame in which it was annotated to occur; this was especially so for death events that were both annotated and predicted. After manual inspections of the mis-predicted events, they appear to fall into a few categories. There were a small number of lifespans in which a censor event was predicted early in the movie, but a death event was annotated to occur later in the movie. Conversely, there were instances of the opposite category. While some of the predictions are incorrect, a majority can be explained by “crowded” catchers, where there are two aging cells near the center of the catcher for a significant number of frames of the time-lapse. Because the annotations contained no information indicating a “cell-of-interest”, the computer vision model and the human annotator could have tracked different cells in the same time-lapse. Therefore it is possible that both the predicted and annotated events are correct and are valid lifespan data.

#### Lifespan Analysis

In order to construct meaningful lifespan data from these two models, their outputs were combined to reconstruct the beginning, the end, the outcome, and the number of divisions each cell was predicted to have been observed. The Kaplan-Meier survival curve generated from these predictions was nearly identical to that generated by human annotation (Fig.3D)(24).

### Lifespan Observations

#### Microdissection Validation of Yeast Lifespan Machine

To validate the Yeast Lifespan Machine platform (both the microfluidic device and automated image analysis pipeline), lifespans generated using this platform and those collected by the traditional micromanipulation assay were compared (4). The Yeast Lifespan Machine was able to produce similar Kaplan-Meier curves to those generated by microdissection for two strains with distinct lifespans (Fig.4A). While the relative lifespan differences between the two strains is very obvious with either of the two methods, there are modest differences between the two types of measurements. The variance between the results could be attributed to differences in the protocols or conditions that the aging cells experience between the two protocols.

**Fig. 4.**
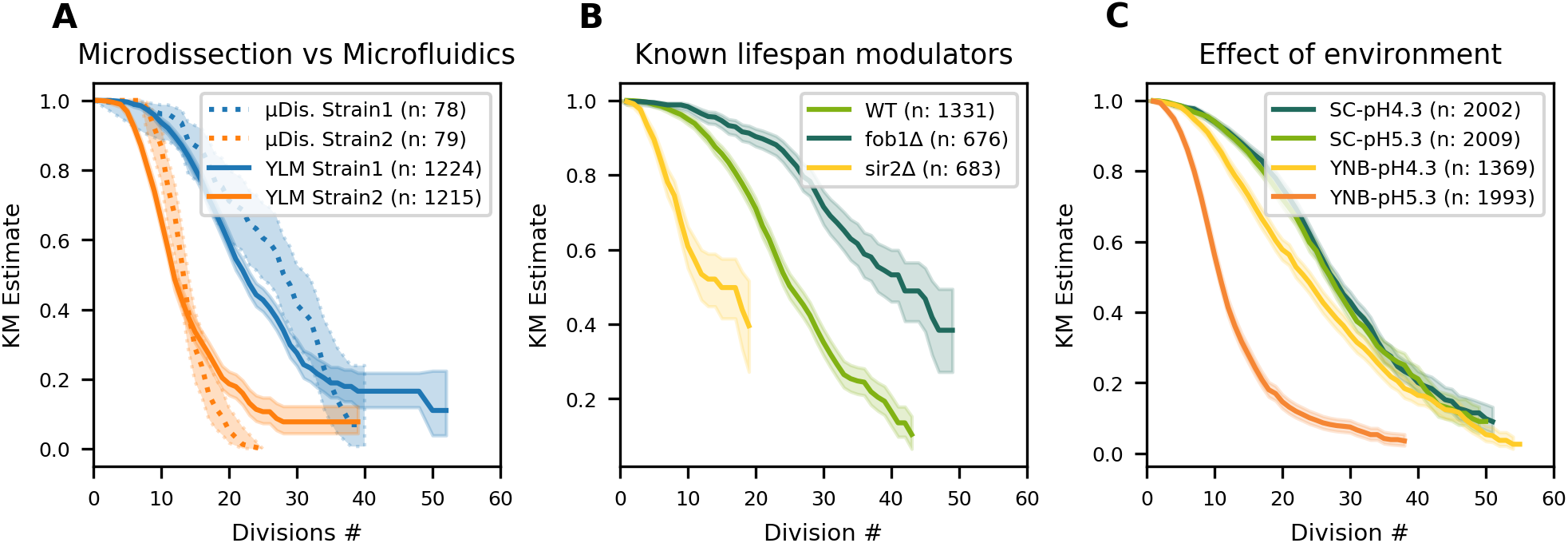
Lifespan observations made with Yeast Lifespan Machine platform. A) Comparison to observations made with traditional micromanipulation assay. Data collected by microdissection is represented by dotted lines; data collected using the Yeast Lifespan Machine (YLM) platform are solid lines. Strains with two different lifespans were measured (blue and orange lines). B) Recapitulation of previously known long- and short-lived mutant strains. The lifespans of *fob1*Δ (long-lived), *sir2*Δ (short-lived) and WT strains were measured with the Yeast Lifespan Machine platform. C) Environmental effects on lifespan. The lifespans of a single strain were measured in multiple different environments (SC: rich synthetic media, YNB: minimal synthetic media; adjusted to pH specified in legend).

#### Previously Known Long- and Short-lived Mutants

As a further evaluation of the Yeast Lifespan Machine’s performance, the RLS of two canonical yeast lifespan modulating mutations was examined: *fob1*Δ and *sir2*Δ. While these two mutations provide a robust increase and decrease, respectively, in lifespan (25, 26), and therefore should be easy for any system to detect, they also present unique tests to our microfluidic strategy and computer-vision pipelines. In addition to altering cell cycle times, *sir2*Δ cells are known to dramatically increase the frequency of elongated daughter morphologies (17), resulting in decreased retention of the *sir2*Δ cells in our device. Nevertheless, the Yeast Lifespan Machine was able to observe a clear reduction in lifespan (Fig.4B). Additionally, a clear increase in the lifespan estimate of *fob1*Δ cells was also observed.

#### Environmental Effects

The environment has long been implicated in modulating the aging process of yeast and other organisms. Microfluidic devices provide a critical tool for investigating these effects because they permit precise control of the cell’s environment throughout the aging process. The constant flow of medium required to prevent the accumulation of daughter cells also blocks the cells’ ability to modify their local environment like they would in batch culture or on an agar plate.

We sought to specifically test the lifespan of cells grown in two nutritional contexts and two environmental pHs. In minimal media (YNB), cells are provided essential vitamins and minerals, a carbon source (glucose) and a nitrogen source (ammonium), and therefore must synthesize all of their own amino acids. In complete media (SC), YNB is supplemented with amino acids and some nucleotides. Due to these differences, there is also a difference in pH. Typically, YNB is pH 5.3 and SC is pH 4.3. However, they are both relatively unbuffered, so their final pH is easily perturbed.

In this experiment, we aged a wildtype prototrophic strain in 4 different environments: i) SC at its unadjusted pH of 4.3 ii) YNB at its unadjusted pH of 5.3 iii) SC adjusted to the pH of YNB (pH 5.3) iv) YNB adjusted to the pH of SC (pH 4.3). Interestingly, while the lifespans of cells aged in SC media were unaffected by changing the pH, the lifespan in YNB was significantly shorter at pH 5.3 (Fig.4C). These results indicate, depending on the environmental context, pH is potentially a significant modulator of lifespan.

## Discussion

In this report, we described both a novel microfluidic device and an automated image analysis pipeline capable of determining a large number of replicative lifespans of budding yeast cells. Our platform represents a significant advance over previously described systems in that it leverages a robust multi-layer valve design to provide a significant increase in the throughput. Combined with the improvements to image analysis, these advances allow us to measure the lifespan under multiple genetic and environmental conditions, all with minimal human setup, intervention, or annotation of the resulting raw data.

We note that a number of other advances in microfluidic technologies and automated image analysis have been applied to this problem and reported by other groups (17, 27, 28). While it is difficult to directly compare these methods, various improvements in microfluidic layout, cell retention features, image collection, and imaging analysis could be combined to create more robust and accessible tools for measuring the lifespan of yeast cells.

While the quantification of RLS using the Yeast Lifespan Machine pipelines is robust within the constraints of the genetic and environmental conditions we tested here, we note that considerable care should be taken when using these or similar microfluidic devices and computer vision models in other contexts. Genetic or environmental conditions that severely perturb the growth rate, morphology, or budding pattern of cells could alter the performance of these systems. It is important that the training data takes into account such features of the data to ensure accurate results. Additionally, the microfluidic catcher features may perform differently in conditions with significantly altered cell morphologies. Perturbations that alter retention rates, especially as cells approach death, could dramatically skew the estimate of the survival function (24).

An exciting aspect of the high-resolution information that we capture using the Yeast Lifespan Machine pipeline is the depth of the time-lapse data, capturing fundamental aspects of yeast aging beyond RLS. Considerable information about the aging process and various aging trajectories are available in these images. Information like cell morphologies, cell division times, and more, could provide significant insights into the mechanisms of aging in this organism (17, 29, 30). Continued development of microfluidic technologies and image analysis pipelines will gain access to this information and improve our understanding of the various aging processes occurring in yeast and ways of modulating them.

In this report, we showed that the Yeast Lifespan Machine platform was able to detect previously reported lifespan modulating effects of four different genetic alterations (all without any further human annotation of the data). In addition, we identified a seemingly subtle environmental change (media pH) that has a strong effect on lifespan. This detail reinforces the importance of controlling quantifiable aspects of the growth environment between experiments and different batches/lots of media in other lifespan studies. Microfluidics provides a valuable tool to investigate relatively subtle changes in the environment, which is simply not possible in batch cultures or cells aging on a plate.

We have only explored a small subset of the possible ranges of physiological tolerable pH ranges and their interactions with different media recipes (31, 32). It is possible that there are pHs that yeast cells are thought to tolerate that may shorten the lifespan in SC or in other environmental contexts. However, we have shown that pH can have a large effect on lifespan, greater than that of many of the genetic perturbations we are investigating. And while the exact mechanism of this effect is not known, at a minimum, the pH of the environment must be considered as a possible source of lifespan variation between conditions, experiments and labs. As these high-throughput lifespan technologies become more widely accessible to the field, we expect that it will be easier to identify and study the sources of variability, potentially reconciling differences in reported lifespan effects of various genetic and environmental perturbations.

## Materials and Methods

### Microfluidic Device

#### Microfluidics Design

Each set of Yeast Lifespan Machine molds were created with four photomasks (Supplemental Material) . One mask is used to pattern positive photoresist (SPR 220-7) onto the fluidics mold to form collapsible channel sections (see below) . Two masks are used to pattern negative photoresist (SU-8) onto the fluidics mold and create channels and catcher features. The last mask is used to pattern negative photoresist onto the control layer mold. All masks containing low resolution features (channels, valves) were printed on transparency masks (Fineline Imaging). Masks with catcher features were printed onto a chrome mask (Compugraphics). Generally we followed the design rules presented by the Stanford Microfluidics Foundry (33). All design work was done in AutoCAD (AutoDesk).

#### Microfluidic Mold Fabrication

The two molds for the Yeast Lifespan Machine device were made using standard multilayer photo-lithography on blank silicon wafers (University Wafer). The fluidics layer mold consists of features created with two types of photoresist: positive and negative. First, the wafer is treated with hexamethyldisilazane vapor for at least 10 minutes. Approximately 2mL of SPR 220-7 (Dow) is spun onto the wafer (500rpm, 1200rpm/s acceleration, 30 seconds; 3000rpm, 1200rpm/s acceleration, 60 seconds) using a spin coater (WS-400-6NPP-LITE, Laurell Technologies, North Wales, PA) and then placed on a hotplate set at 90° C (Torry Pines HP40A, Torrey Pines, Carlsbad, CA) for 90 seconds. The valve pattern is then transferred to the wafer using a contact aligner (Karl Suss Ma6, Suss MicroTec, Garching Germany) with LED illumination in 37% i-line only mode using Proximity Exposure at 30 μm and an exposure of 380 mJ/cm^2^. The wafer is then developed manually in MF26A developer (Dow) and then hard-baked on a hotplate to melt and reflow the photoresist starting at room temperature and then ramping up to 190° C at 15° per hour for a total of 20 hours. After hard baking the valves are measured using a KLA D-600 profilometer (KLA, Milpitas, CA) and the rounded features are around 4.5 μm tall. A 6.5 μm layer of SU-8 2005 (Kayaku Advanced Materials, Westbough MA) is spun on the wafer (500rpm, 1200rpm/s acceleration, 30 seconds; 1050rpm, 1200rpm/s acceleration, 30 seconds). Then the wafer is baked on a hotplate at 65° C for 3 minutes and then at 95° C for 5 minutes. The catcher pattern is transferred to the wafer using a contact aligner and an exposure of 130-140 mJ/cm^2^. Then the mask is changed to expose the remaining channel features in the fluidics layer, using Proximity Exposure at 30 μm and an exposure of 300 mJ/cm^2^. Then the wafer is baked on a hotplate at 65° C for 3 minutes then at 95° C for 5 or 6 minutes. After baking the wafer it is dipped in Su8 developer (Kayaku Advanced Materials, Westbough MA) and agitated for 20-40 seconds until the non-crosslinked Su8 is removed. The wafer is then sprayed with isopropyl alcohol and air dried with a nitrogen gun. After drying the wafer is inspected for signs of residual uncrosslinked SU-8 such as white streaks on the wafer or scumming around the features. If needed, the wafer is dipped in the developer for another 5-10 seconds, sprayed with isopropanol, and air dried. The control layer mold was made with negative photoresist using a similar protocol as outlined above with the following modifications: Feature height ~25*μ*m; SU-8 2025 spun on at 2100 rpm; exposure 150 mJ/cm^2^; Post Exposure Bake 1 65° C for 1 minute then at 95° C for 3 minutes. Specific spin, bake and exposure parameters may require additional optimization on other equipment.

#### Microfluidic Device Fabrication

All mixing, spinning, and room temperature incubations are carried out in a humidity controlled chamber, held at 15% RH. This is critical for reliable operation of the microfluidic valves. First, PDMS (Momentive #RT615; 5 part A: 1 part B) is poured onto control layer mold, degassed under vacuum for 40 minutes, then baked at 80° C for 60 minutes. Second, a thin layer of PDMS (20 part A : 1 part B) is spun coat onto fluidic layer mold (1.25 minutes at 900 RPM) and baked at 80C for 40 minutes. The control layer is then removed from mold, cut to size, and holes are punched to access valve lines. This PDMS block is then aligned to the fluidics layer under a microscope. The aligned device is then baked at 80° C for an additional >4 hours. Devices are then removed from fluidic layer mold, access ports punched, and then plasma bonded to a large glass coverslip.

### Experiment Acquisition/Set-Up

### Strains, Media, Culturing

All strains presented in this study and used for the optimization of the Yeast Lifespan Machine chip are haploid strains derived from the BY strain background. See Table 1.

Note, CGY2.66 has a Ty element inserted into *PTR3*. While the function of *PTR3* in this strain has not been investigated, other experiments in our lab have shown that this polymorphism is not responsible for lifespan differences in media of different pH.

**Table 1.** Strain Table.

| Name | Genotype | Source | Figure |
| --- | --- | --- | --- |
| CGY20.46 | *MATa can1Δ::STE2pr-spHIS5 his3Δ1 lyp1Δ0* | Hotz et al. 2022 / This Study | 2 |
| CGY20.50 | *MATa can1Δ::STE2pr-spHIS5 his3Δ1 lyp1Δ0* | Hotz et al. 2022 / This Study | 2 |
| CGY20.55 | *MATa can1Δ::STE2pr-spHIS5 his3Δ1 lyp1Δ0* | Hotz et al. 2022 / This Study | 2 |
| CGY24.09 | *MATa can1Δ::STE2pr-spHIS5 his3Δ1 lyp1Δ0* | Hotz et al. 2022 / This Study | 2 |
| CGY24.16 | *MATa can1Δ::STE2pr-spHIS5 his3Δ1 lyp1Δ0* | Hotz et al. 2022 / This Study | 2 |
| CGY24.17 | *MATa can1Δ::STE2pr-spHIS5 his3Δ1 lyp1Δ0* | Hotz et al. 2022 / This Study | 2 |
| CGY24.21 | *MATa can1Δ::STE2pr-spHIS5 his3Δ1 lyp1Δ0* | Hotz et al. 2022 / This Study | 2 |
| CGY28.49 | *MATa can1Δ::STE2pr-spHIS5 his3Δ1 lyp1Δ0* | Hotz et al. 2022 / This Study | 2 |
| CGY28.50 | *MATa can1Δ::STE2pr-spHIS5 his3Δ1 lyp1Δ0* | Hotz et al. 2022 / This Study | 2 |
| CGY28.52 | *MATa can1Δ::STE2pr-spHIS5 his3Δ1 lyp1Δ0* | Hotz et al. 2022 / This Study | 2 |
| CGY28.53 | *MATa can1Δ::STE2pr-spHIS5 his3Δ1 lyp1Δ0* | Hotz et al. 2022 / This Study | 2 |
| CGY29.01 | *MATa can1Δ::STE2pr-spHIS5 his3Δ1 lyp1Δ0* | Hotz et al. 2022 / This Study | 2 |
| CGY24.09 | *MATa can1Δ::STE2pr-spHIS5 his3Δ1 lyp1Δ0* | Hotz et al. 2022 / This Study | 4A |
| CGY24.17 | *MATa can1Δ::STE2pr-spHIS5 his3Δ1 lyp1Δ0* | Hotz et al. 2022 / This Study | 4A |
| CGY20.74 | *MATa* | This Study | 4B |
| CGY19.41 | *MATa fob1Δ::NatMX* | This Study | 4B |
| CGY20.67 | *MATa sir2Δ::KanMX* | This Study | 4B |
| CGY2.66 | *MATa ptr3::Ty* | This Study | 4C |

**Table 2.** Parts List. Key materials to assemble acquisition hardware and microfluidic devices.

|  | Part Name | Supplier |
| --- | --- | --- |
| Microscope Stand | RAMM system equipped with MS-2000 automated XY-stage | Applied Scientific Instrumentation |
|  | Transillumination kit based | Applied Scientific Instrumentation |
|  | pco.panda 4.2 sCMOS camera | PCO |
|  | CFI60 Plan Apo Lambda 40x Objective Lens, .95NA | Nikon Instruments Inc. |
|  | Incubator assembled from laser-cut acrylic and optomechanical components | Custom (supp. mat., Thorlabs) |
|  | Multiwell microplate insert | Applied Scientific Instrumentation |
| Microfluidic Controls/Accessories | Device holder | Custom-machined (supp. mat.) |
|  | Microfluidic valve controller | Elveflow or Custom-built |
|  | Pressure regulating flow controller | Elveflow or Custom-built |
|  | Compression fittings - Tuohy Borst 1032, 1/32"-1/16", 3fr-5fr | Microgroup Inc. |
|  | Custom manifold for 24-well plate | Custom-machined (supp. mat.) |
|  | Custom manifolds for media reservoirs | Custom-machined (supp. mat.) |
|  | Pressure regulator flow controller | Elveflow or Custom-built |
| Consumables | Fluidic tubing - PTFE tubing ID:0.022±0.002" OD:0.034" | Component Supply |
|  | Device adapter pins - 304/306 SS Tubing OD:0.025"xID:0.013"xL:0.75" | New England Small Tube Corp. |
|  | Reservoir "spear" pin - 304/306 SS Tubing OD:0.025"xID:0.013"xL:6" | New England Small Tube Corp. |

Experiments were conducted in either i) YPD – 1% w/v Yeast extract, 2% w/v Bacto peptone and 2% glucose ii) Synthetic Complete (SC) media – 1x SC mix and 1x YNB with Ammonium Sulfate (Sunrise Science), 2% glucose iii) Minimal media (YNB) – 1x YNB with Ammonium Sulfate (Sunrise Science), 2% glucose. pH was adjusted as noted using Potassium Hydroxide.

Experiment presented in machine learning validation (Fig. 3) was conducted in SC. Experiment presented in comparison to micro-dissection was conducted in YPD (Fig.4A). Experiment measuring the lifespans of *sir2*Δ and *fob1*Δ strains was conducted in SC (4B). Experiment exploring the effect of pH in different nutrient conditions (4C) was conducted as described. Rhodamine labeling of original cells was achieved by incubating cells in NHS-Rhodamine (1ug/ml; Fisher Scientific) for 5 minutes before loading into the microfluidics device (34).

To begin a lifespan experiment, cells were grown to saturation in 24-deep-well plates, then diluted in fresh media and grown for 24 hrs below an OD of 0.05. Cells were then loaded into the microfluidics device by pneumatic pressure using a custom machined manifold (supplemental materials). All experiments were conducted at 30° C.

Microdissection RLS measurements were performed similar to previously described (4). Cells were grown in liquid YPD before being spotted onto YPD agar plate. Cells were then moved using a microdissection needle (Singer Instruments) to a unique position on the plate. After these cells completed their first division, the newly born cell was kept to begin the lifespan measurement. After each division, daughter cells were removed until the mother cell stopped dividing or the experiment was terminated. Any mother cell that was still dividing at the end of the experiment was considered censored (see Lifespan Analysis methods section).

#### Image Acquisition Hardware

A parts list of our custom assembled microscope is available in the supplemental material (2. A majority of our data were collected on an ASI RAMM system equipped with an ASI MS-2000 automated XY-stage, pco.panda 4.2 sCMOS camera, bright-field transillumination kit, and 40x air objective (Nikon NA .95; MRD00405). Illumination and camera triggering were achieved through an analog out card (National Instruments). Environmental control (temperature) was achieved by a custom designed and assembled chamber (supplemental material) and commercially available temperature controller (Oko Labs). Visualization of rhodamine labeled original cells was a Leica DMI8 microscope (Leica Microsystems) equipped with a 40x air objective, pco.Edge 4.2, and a spectraX LED lightsource (Lumencore).

#### Microfluidics Hardware

Microfluidic valves were controlled by computer operated pneumatic solenoid valves (Elveflow or custom assembled controller (35)). Media was driven from 50ml conical tubes (1 per channel) into the device by constant pneumatic pressure provided through custom manifolds (supplemental materials). Pressure was regulated by computer controlled regulators, and optimized to deliver approximately 5μl/minute (Elveflow or custom assembled). PTFE tubing connected media reservoirs to microfluidic devices. Bent metal pins/tubes connected PTFE tubing into PDMS microfluidics devices. Longer metal pins/tubes served as a “spear” to access media in each reservoir.

#### Experiment Acquisition Software

Almost all hardware was controlled with python using custom device adapters (available here: https://github.com/AndrewGYork/tools). Leica instrument was controlled with python through micro-manager and the python bindings for MMCore (micro-manager/pymmcore - Python bindings for MMCore, (36)). Experiment schedule (image frequency/timing) and meta-data were stored in a centralized database. Experiment progress was monitored through communication to the central database and monitoring scripts running on a central server. User notifications of progress, alerts, failures were accomplished with the Google Chat API.

### Analysis Pipeline

#### Image Processing

All image processing steps were completed in python. Raw image time series were aligned with code available in https://github.com/AndrewGYork/tools. Each catcher location was identified by convolving an image with an image of a template catcher and finding local maxima. 256×256 pixel patches were cropped around each identified catcher location. The image patches were used as inputs for subsequent computer vision steps.

#### Computer Vision Models

##### Training Data Collection

Most human annotations were collected using a custom written annotation software built from the napari python package (37). Specific images to annotate were randomly selected and presented to users (blinded to environment and genetic metadata).

#### Division Detection Model and End-Event Detection Model

The models were trained using a Pytorch framework (38). Adam optimizer (39) was used with beta1=0.9 and beta2=0.8, and a step learning rate schedule as follows:

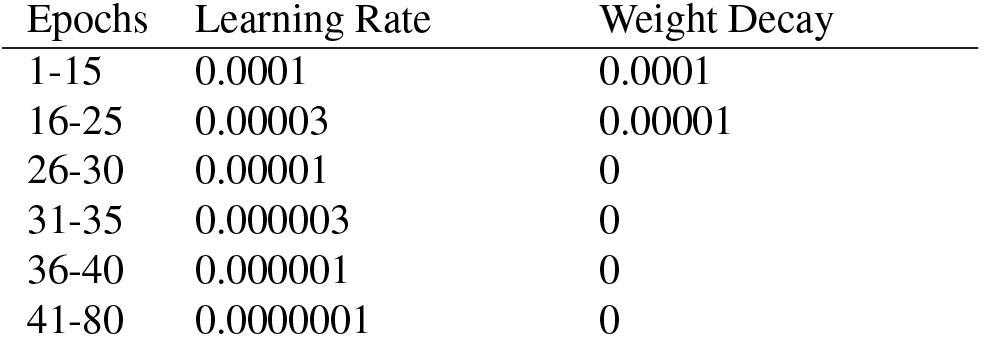

#### Lifespan Analysis

Lifespans were constructed by summing the number of division events that were predicted prior to each predicted-end event. Any divisions that were predicted after an end-event were attributed to a new cell/lifespan. Any lifespan that began after frame 50 (12 hours) was excluded from future analysis. Survival functions were estimated using the Kaplan-Meier estimator, implemented in the lifelines python package (40). Cells that were predicted to be censored by the computer vision model were “right-censored” in Kaplain-Meier fitter (’ end-event=False’ ).

#### Code Availability

Computer vision models are available at https://github.com/calico/ylmcv. Code used to control microscope hardware and steps in image processing steps in https://github.com/AndrewGYork/tools.

## Supporting information

Designs and Schematics

## AUTHOR CONTRIBUTIONS

NHT, MR, JX, ELS, RSM, DEG conceived of the experiments and processes. NHT, MR, ELS, RK, AMS, AGY, AJW developed acquisition instrument and infrastructure. NHT, MR, ELS, MH, JVR, AJW, BW collected data. JX developed computer vision models. NHT, MR, JX, ELS, MH, VO, AJW, RSM, DEG prepared manuscript. MR is presently at Deepcell Inc., Menlo Park, CA 94025, USA. RSM is presently at Foresite Capital, Larkspur, CA 94939, USA. There are no competing interests.

## ACKNOWLEDGEMENTS

We would like to thank: members of the Gottschling lab, Anastasia Baryshnikova, and David Botstein for critical reading of the manuscript. Mike Ando, Benjamin Cooper, Chiraj Dalal, Archa Jain, Meng Jin, and Eddie Xue for technical expertise and various contributions. Alex Chekholko for advice, expertise and maintenance of computing resources.

## Supplementary Note 1: Supplemental Figures and Tables

**Supplementary Figure 1.**
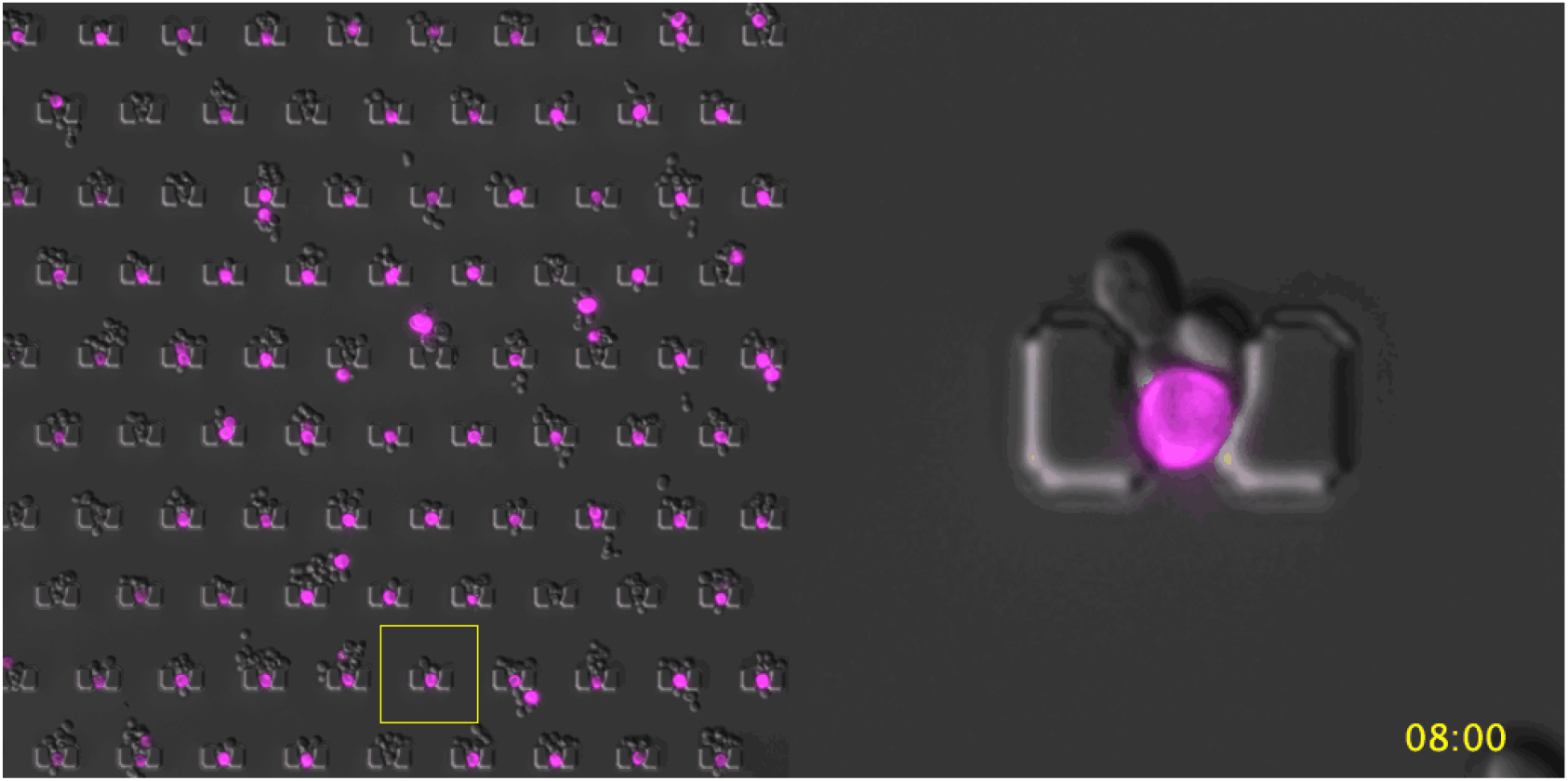
Aging Cell Retention. Original cells were permanently labeled with NHS-Rhodamine prior to loading into the microfluidic device. Any daughter cells produced in the device are unlabeled. The inset highlights a catcher that performs as designed (e.g. retention of original cell until death, washing away of daughter cells, minimal crowding). However, there are a number of other catchers that highlight other situations: i) washout/escape of original cell ii) prolonged retention of multiple cells before they are washed away iii) dramatically altered cell morphologies iv) crowding/clogging of regions of the device.

**Supplementary Figure 2.**
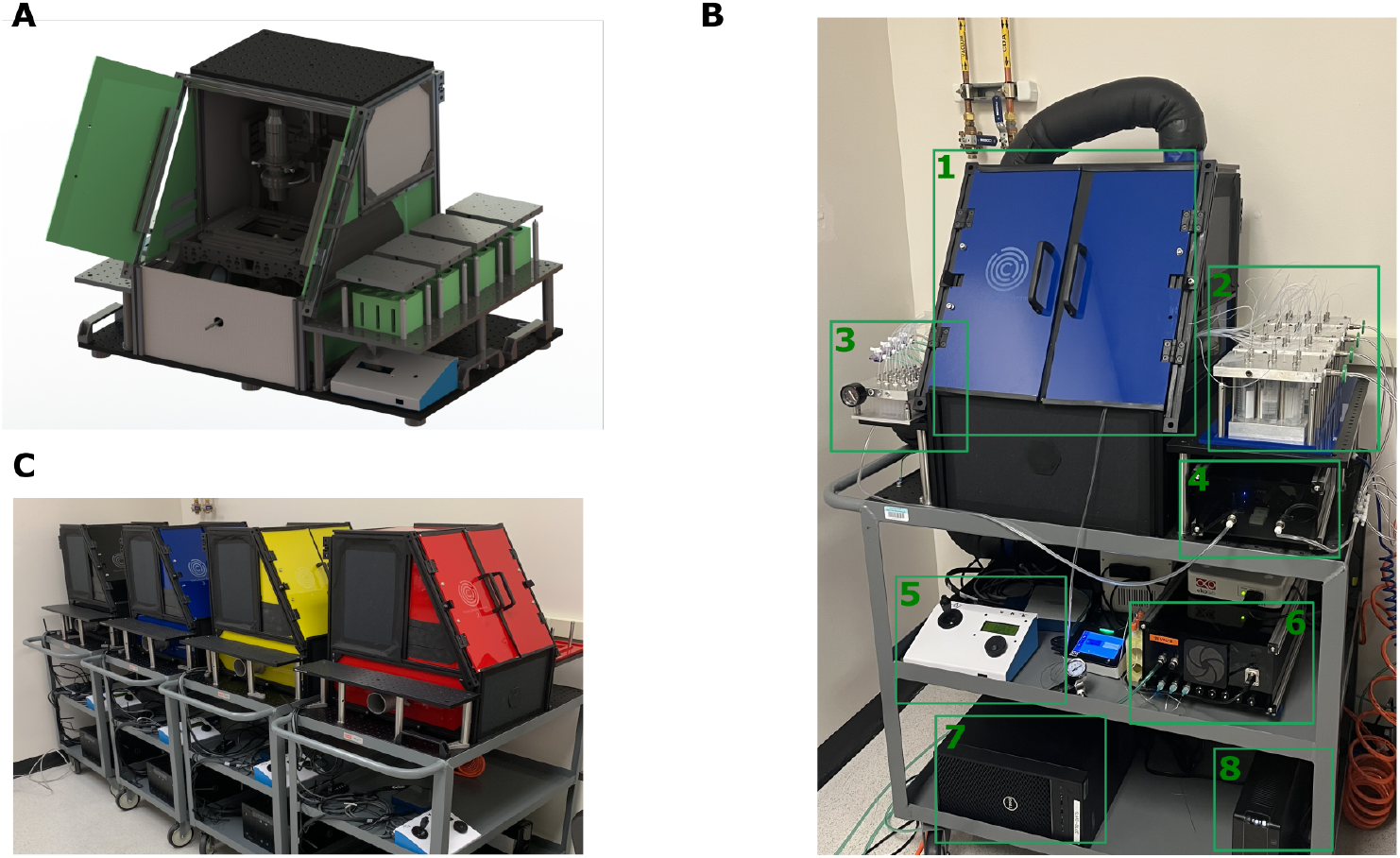
Yeast Lifespan Machine instrument. A) CAD rendering of instrument. Layout of instrument, incubation chamber, and associated microfluidics peripherals (reservoirs, etc) was optimized to reduce footprint and lengths of tubing. B) Image of assembled Yeast Lifespan Machine instrument and accessories. Boxes highlight various components: 1 - microscope and incubation chamber. 2 - media reservoirs and manifolds. 3 - 24-well plate manifold used for cell loading. 4 - computer controlled regulator for managing flow-rates. 5 - the stage controller. 6 - microfluidic valve controller. 7 - the computer used to control each instrument. 8 - uninterruptible power supply, capable of powering the entire instrument during intermittent power outages. C) Image of several instances of Yeast Lifespan Machine Instrument. Optimizations of cost and reliability of the hardware allow us to have up to five instruments running simultaneously (four pictured).

**Supplementary Figure 3.**
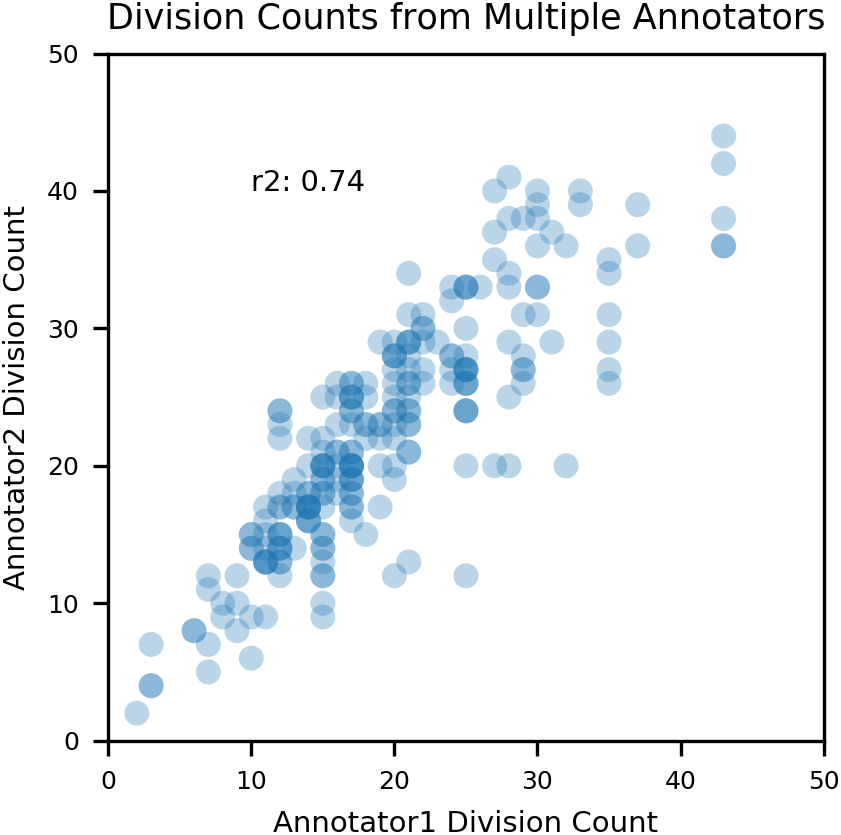
Comparing Annotations From Multiple Annotators. Comparing the total number of annotated divisions from lifespans in catcher time-lapses that had been annotated by two different annotators.

