## Supplementary figures and images for "The Yeast Lifespan Machine: a microfluidic platform for automated replicative lifespan measurements"

### Basic lifespan machine incubator door 1.0 - cut sheet.PDF

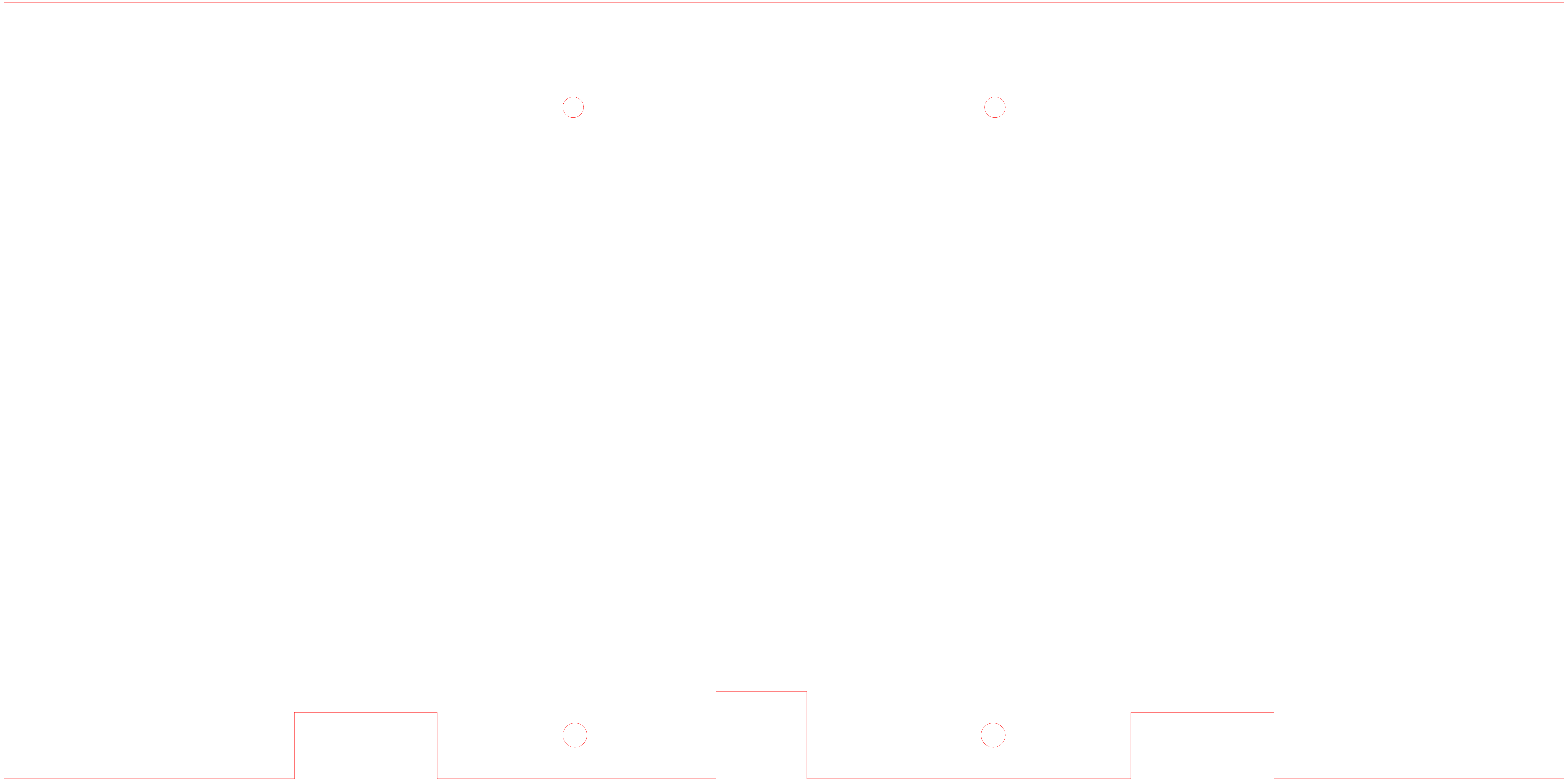

### Basic lifespan machine incubator front lower panel 1.0 - cut sheet.PDF

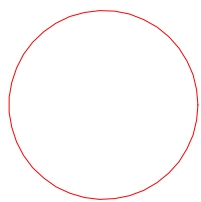

### Basic lifespan machine incubator front lower panel rubber seal 1.0.PDF

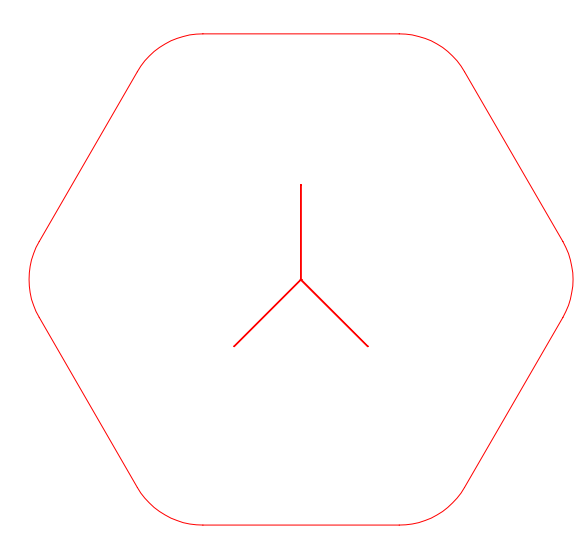

### Basic lifespan machine incubator lower side panel 1.0 - cut sheet.PDF

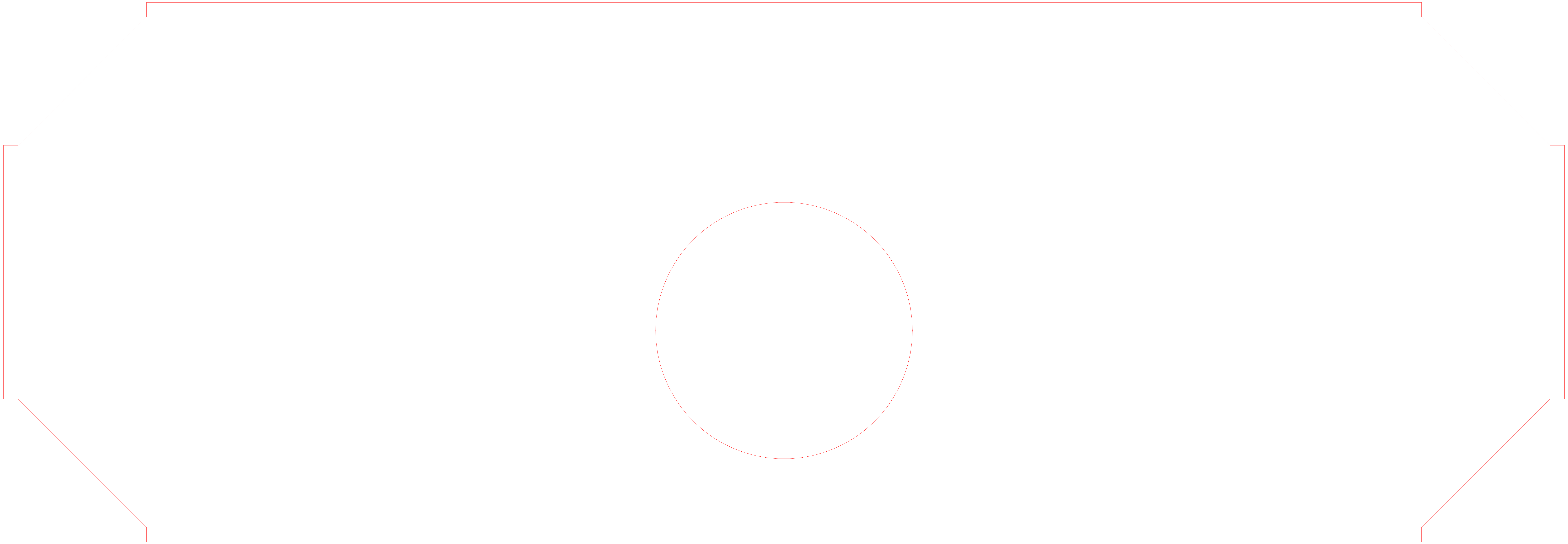

### Basic lifespan machine incubator lower side panel rubber seal 1.0 - cut sheet.PDF

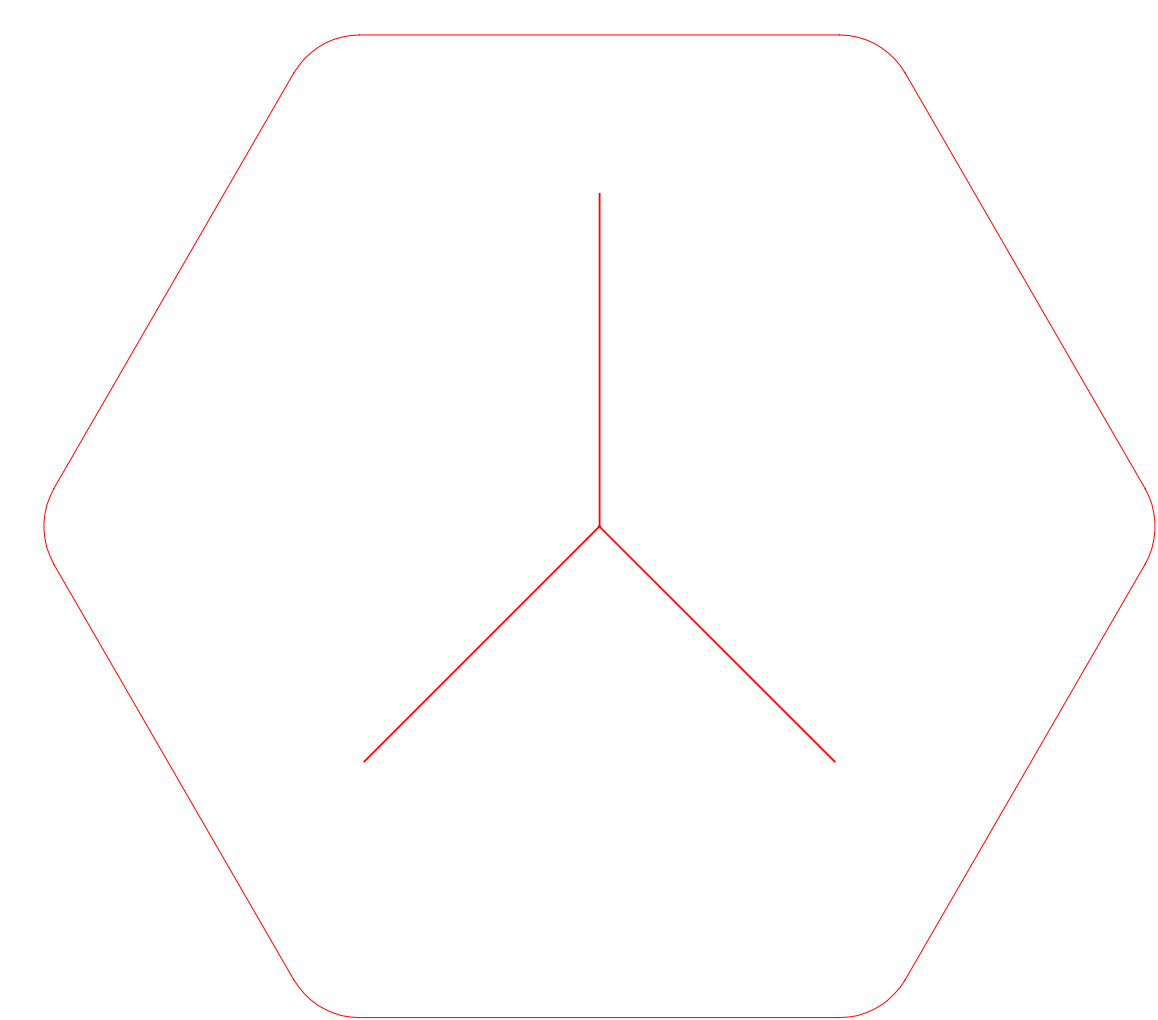

### Basic lifespan machine incubator rear panel 1.0 -- cut sheet.PDF

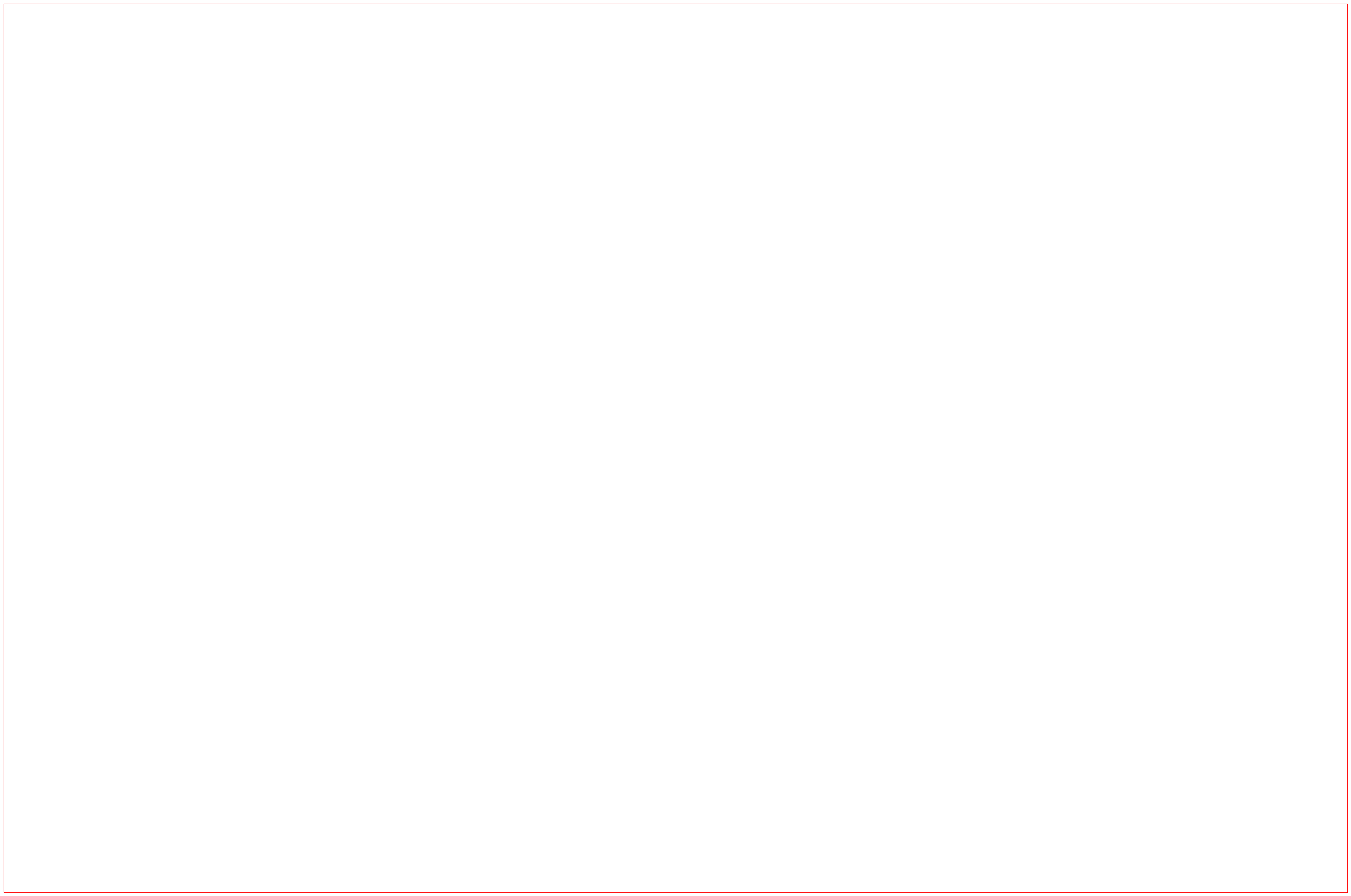

### Basic lifespan machine incubator rear panel 1.0.pdf

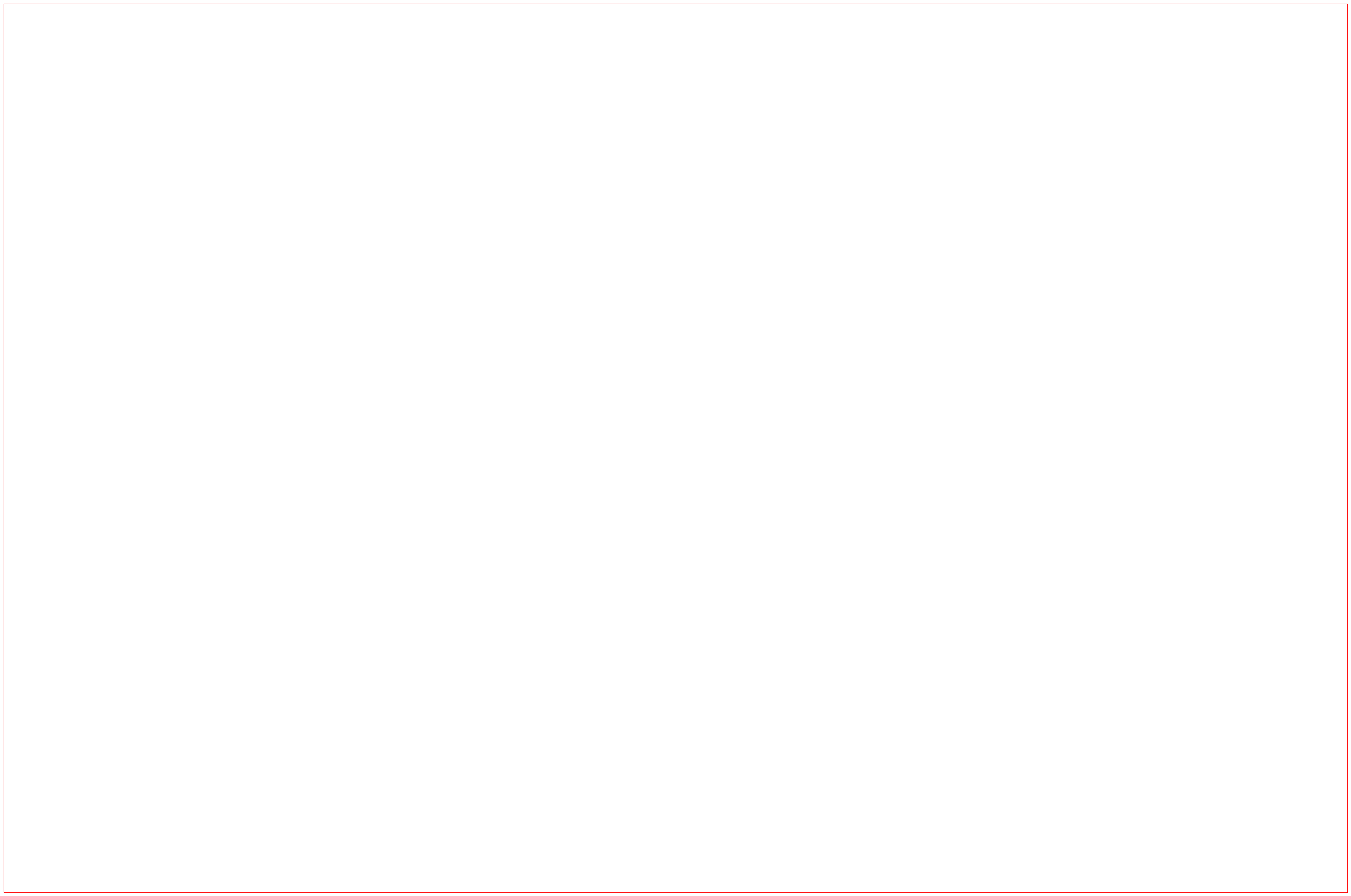

### Basic lifespan machine incubator top panel 1.0 - cut sheet.PDF

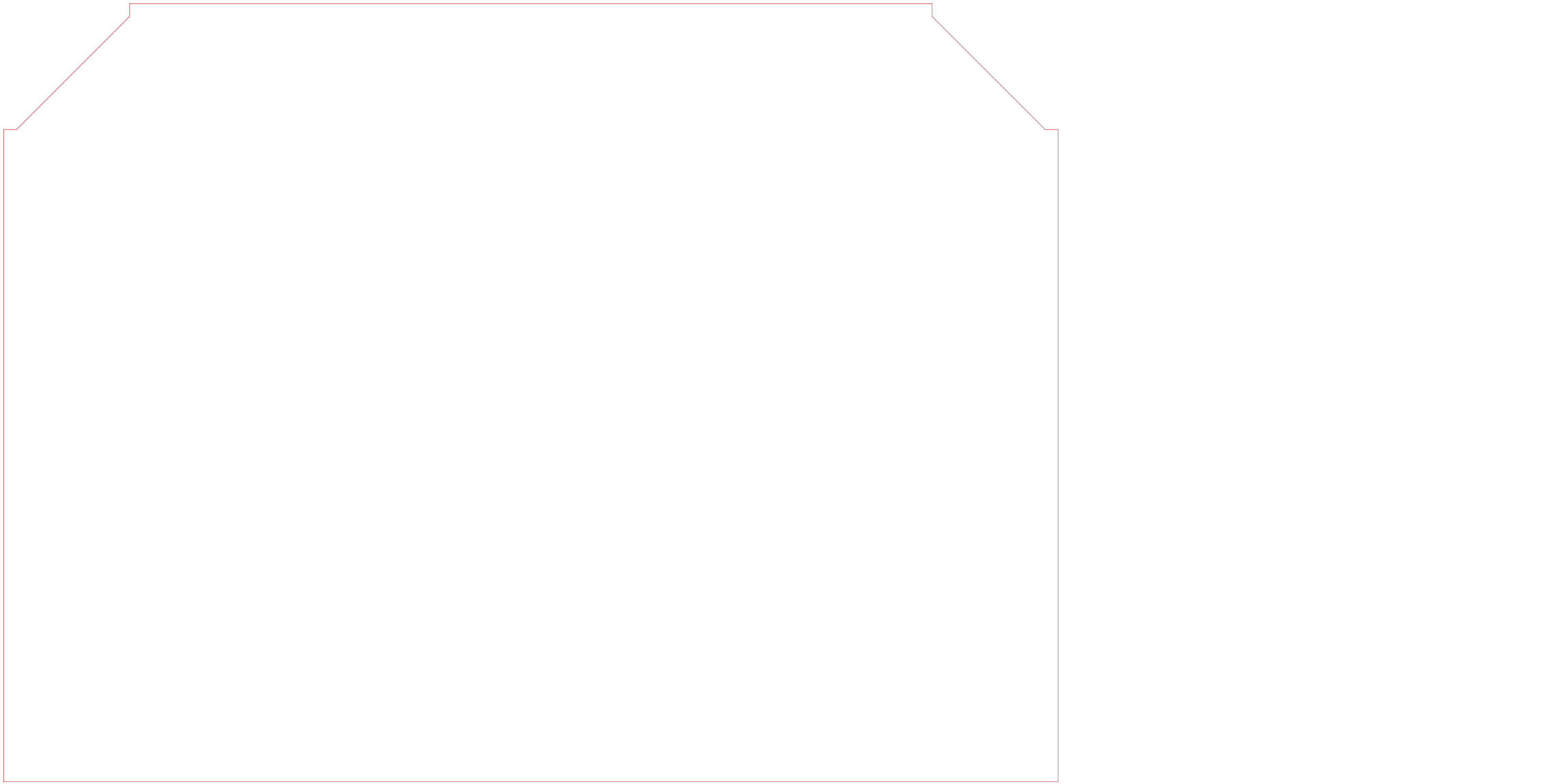

### Basic lifespan machine incubator top panel - with duct port 1.0.PDF

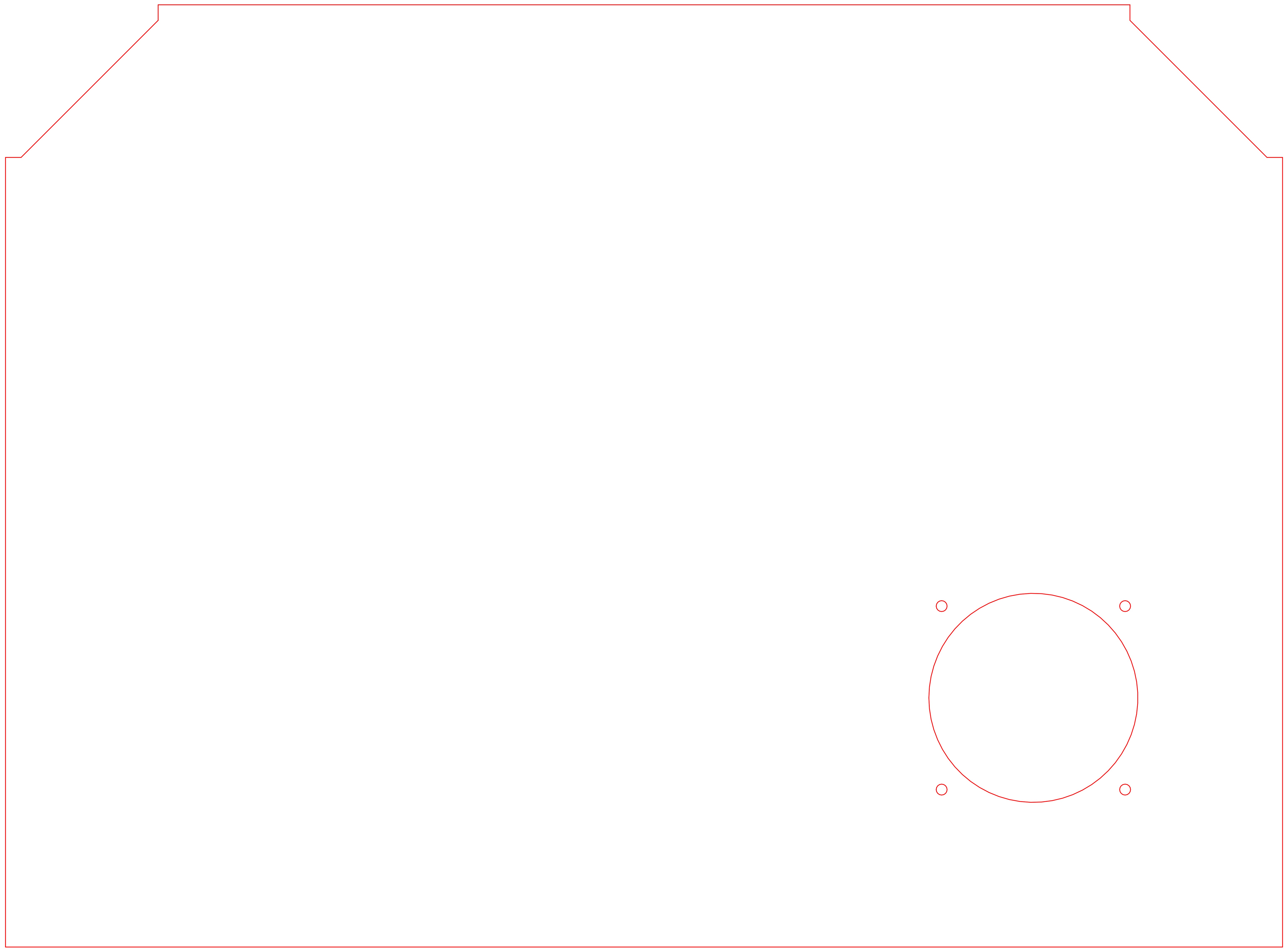

### Basic lifespan machine incubator upper front side panel 1.0 - cut sheet.PDF

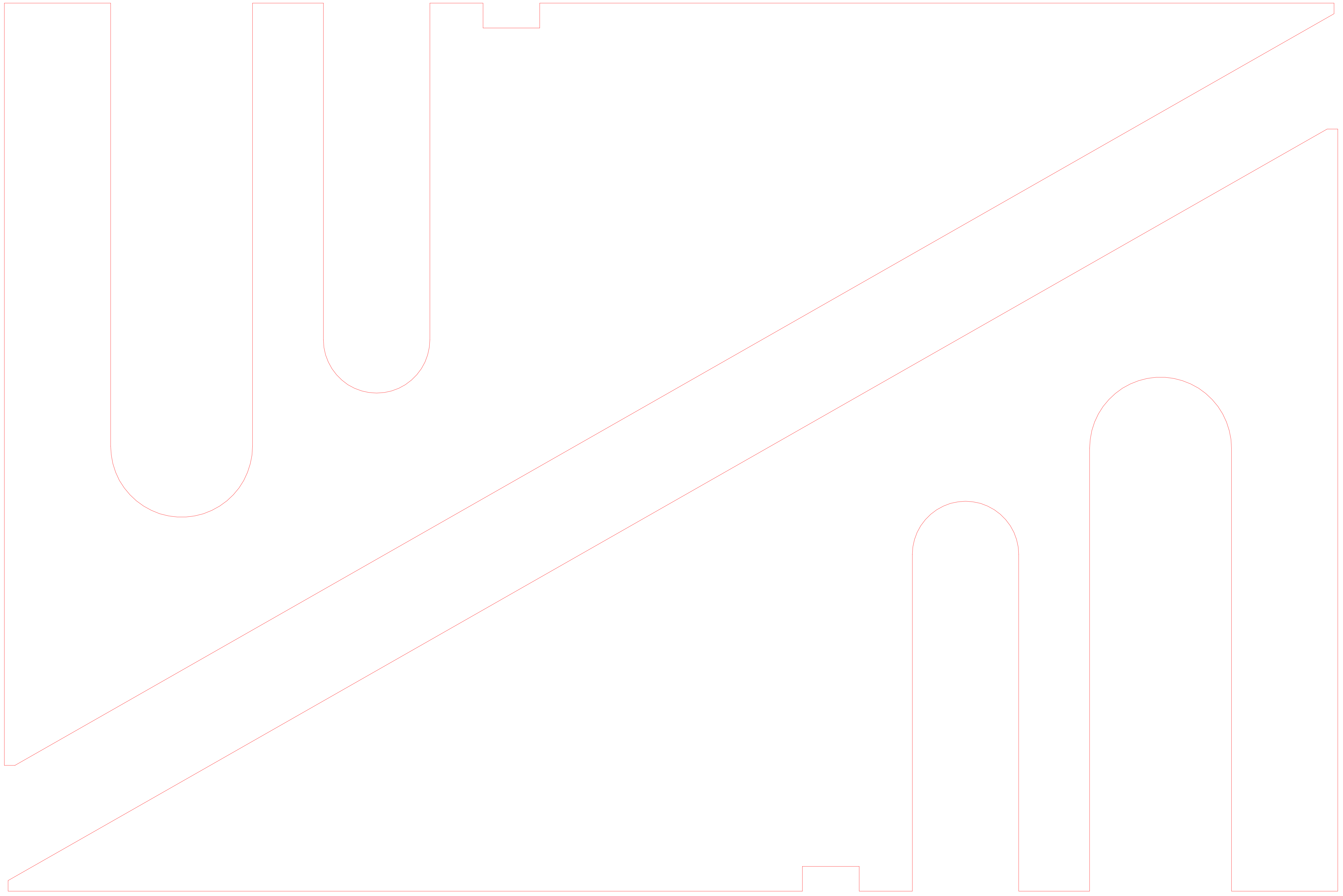

### Basic lifespan machine incubator upper left side panel rubber seal 1.0 - cut sheet.PDF

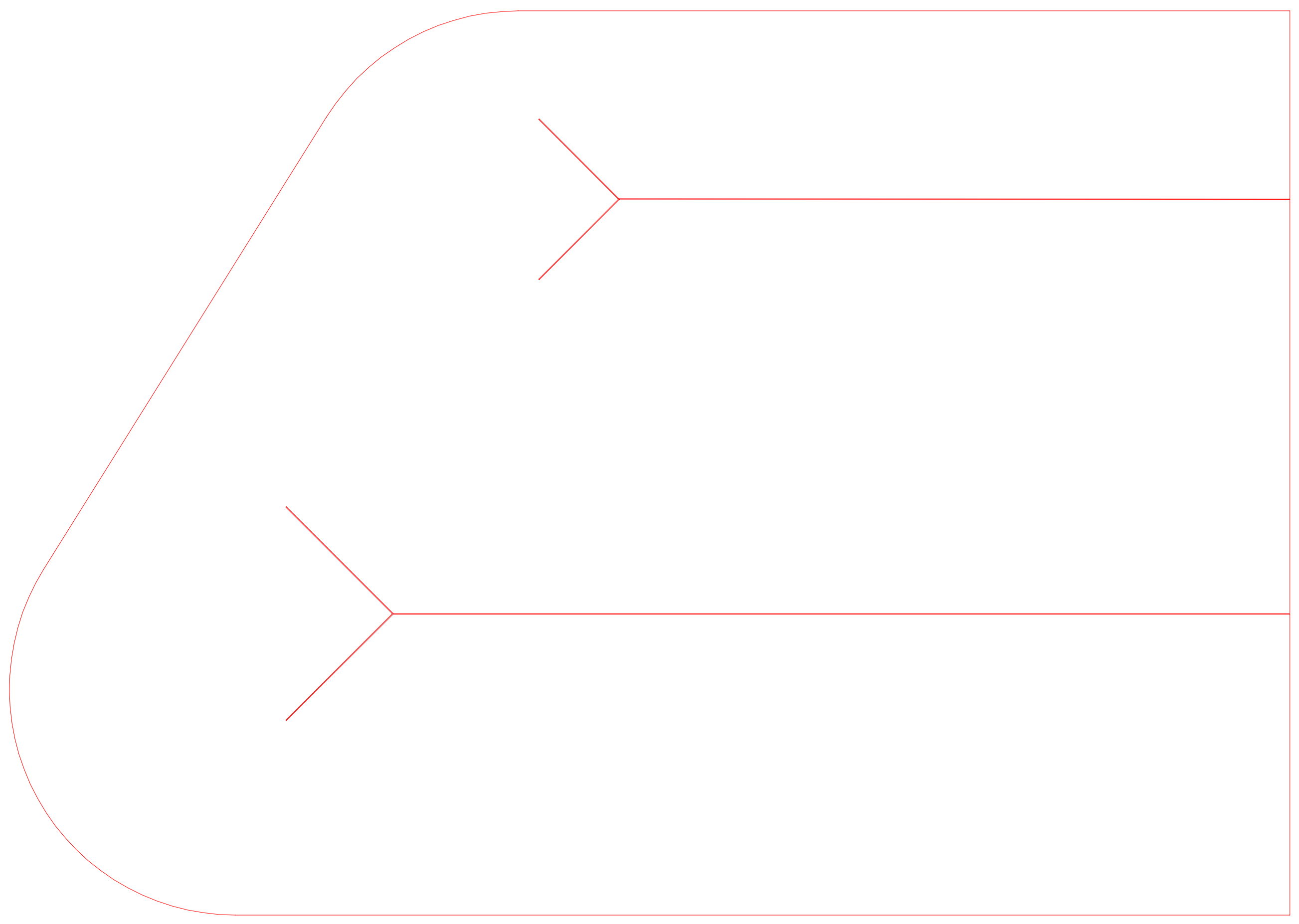

### Basic lifespan machine incubator upper rear left side panel 1.0 - cut sheet.PDF

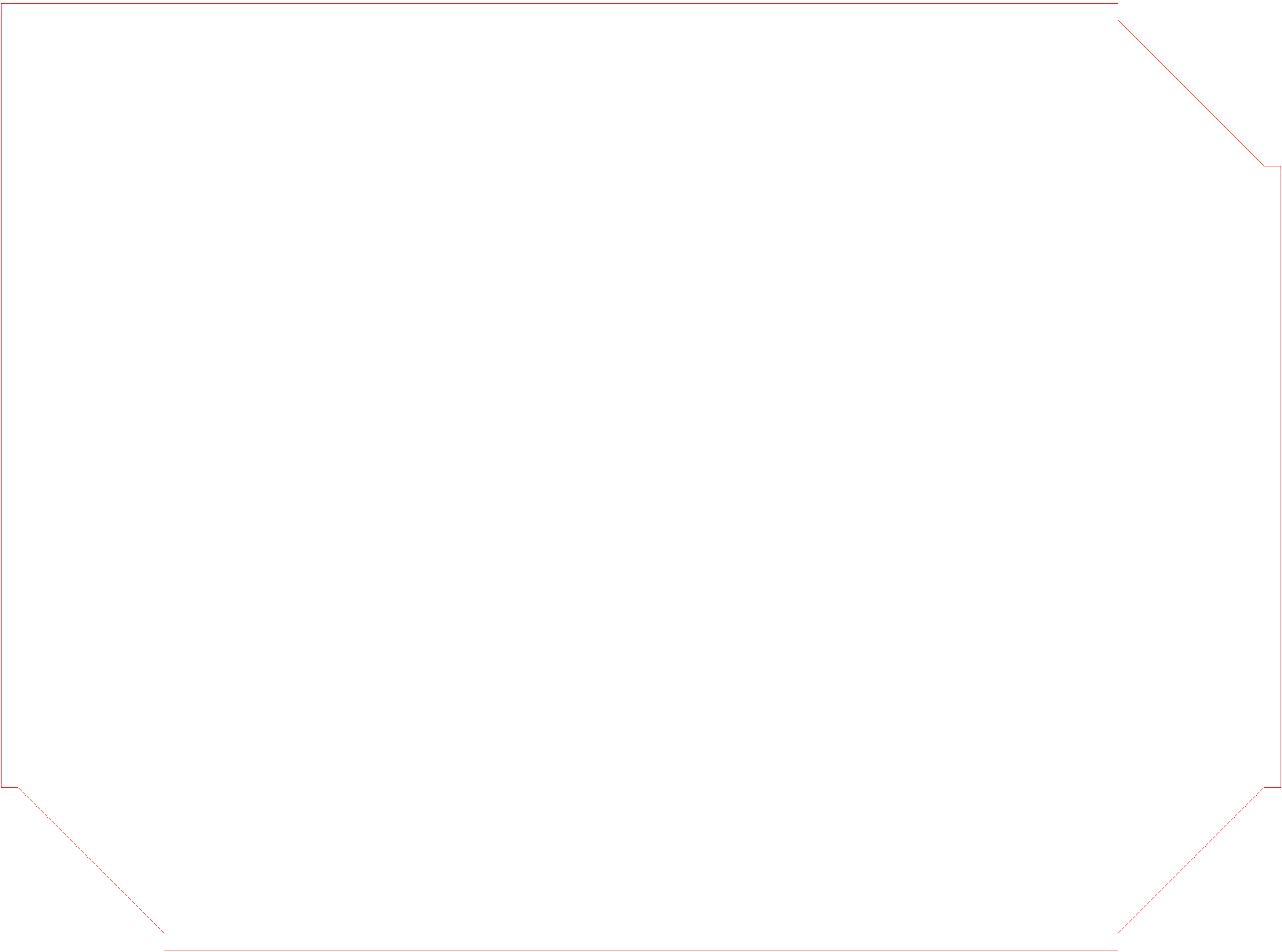

### Basic lifespan machine incubator upper rear right large side panel 1.0 - cut sheet.PDF

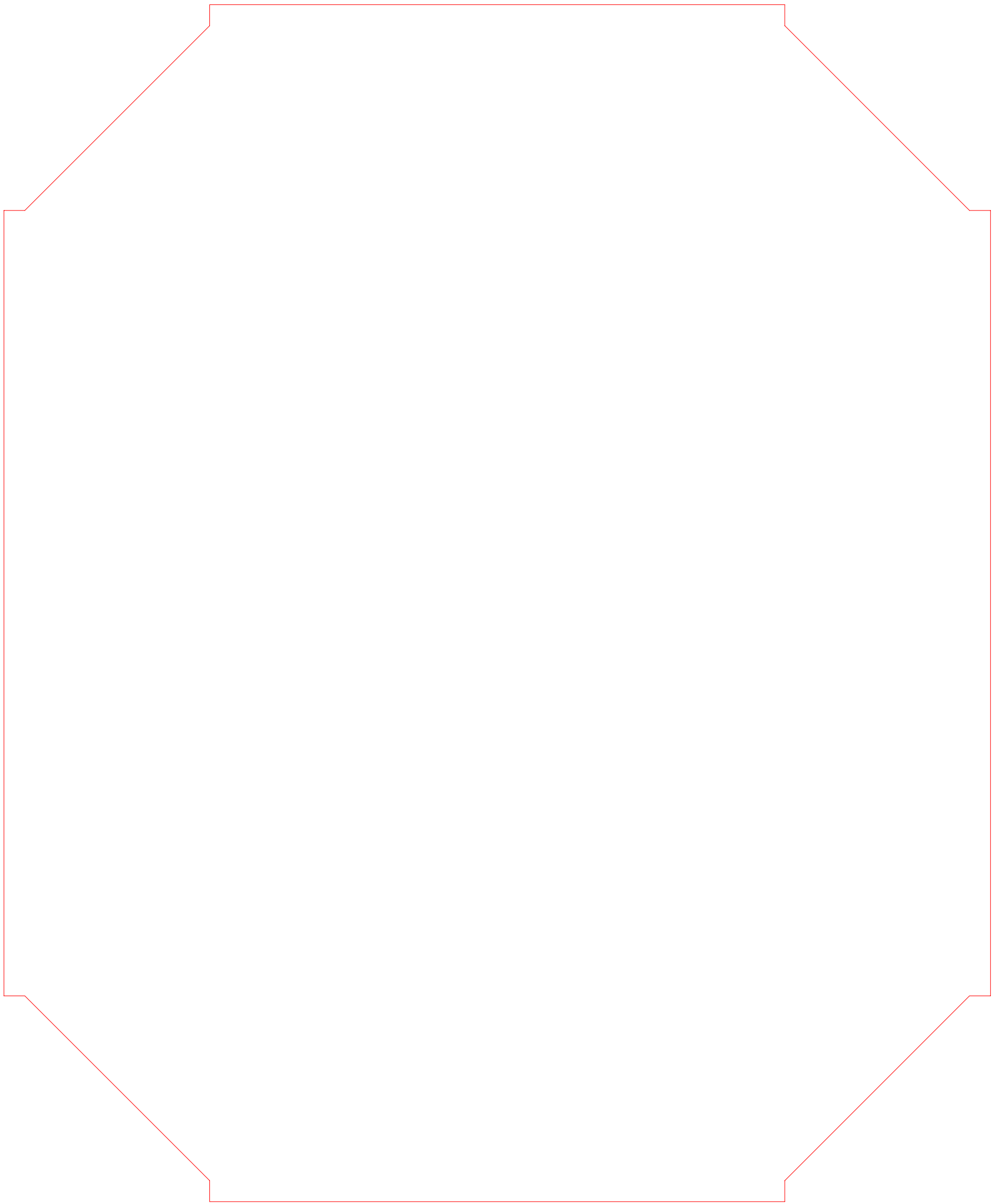

### Basic lifespan machine incubator upper rear right small side panel 1.0- cut sheet.PDF

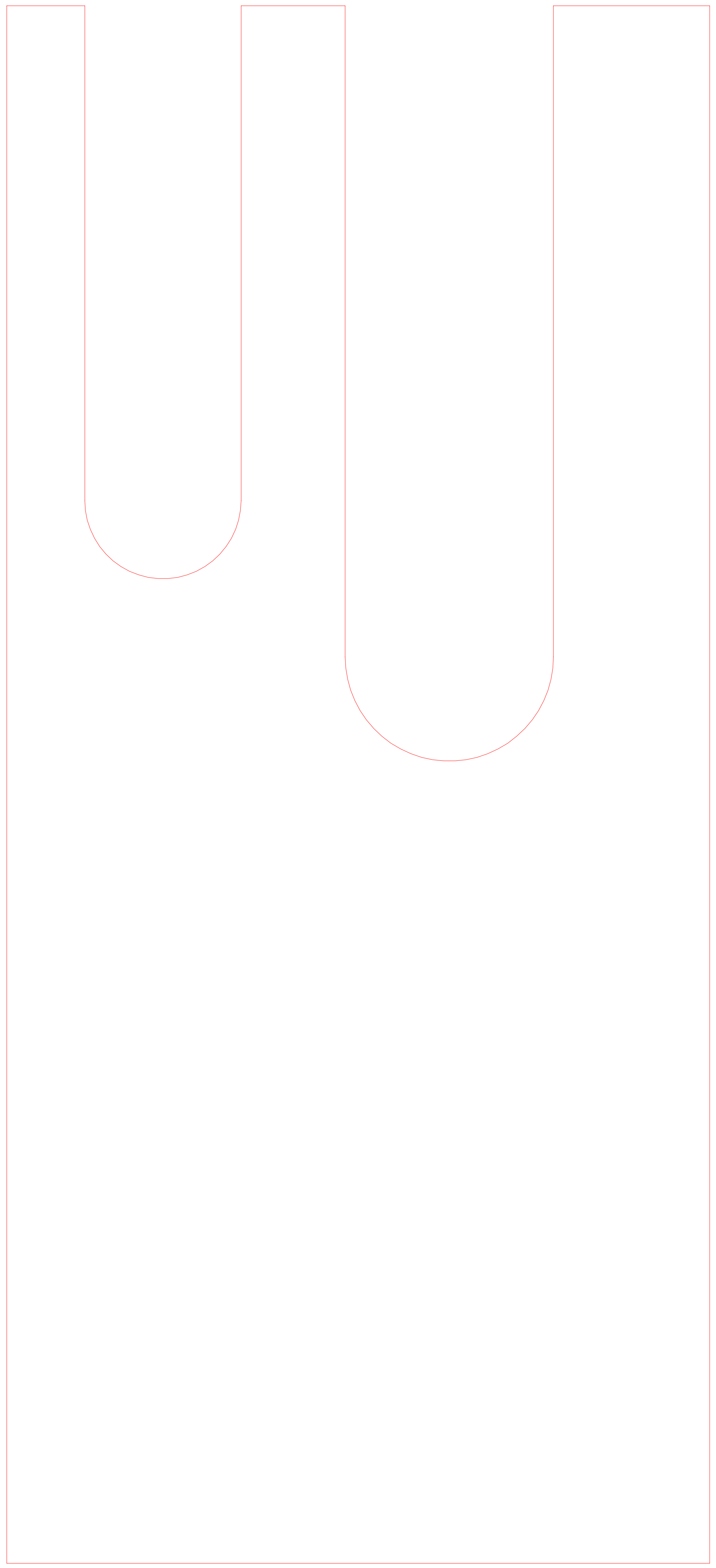

### Basic lifespan machine incubator upper right side panel rubber seal 1.0 - cut sheet.PDF

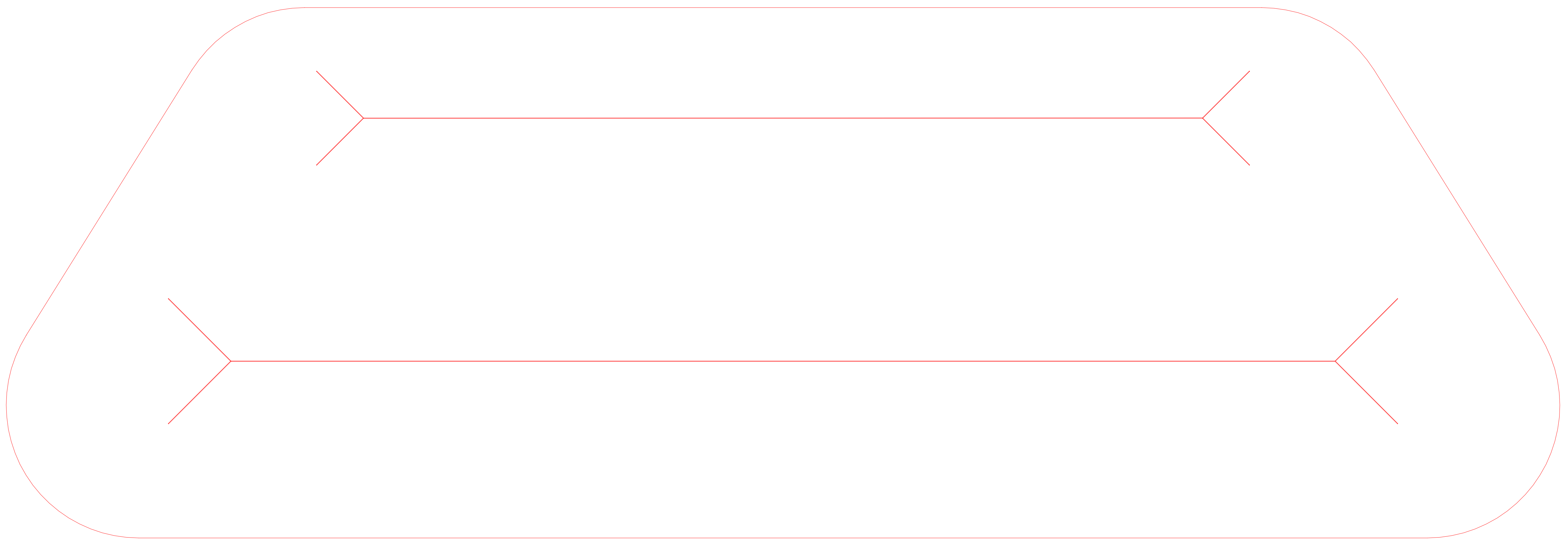

### Basic lifespan machine media tray base 1.0 - cut sheet.PDF

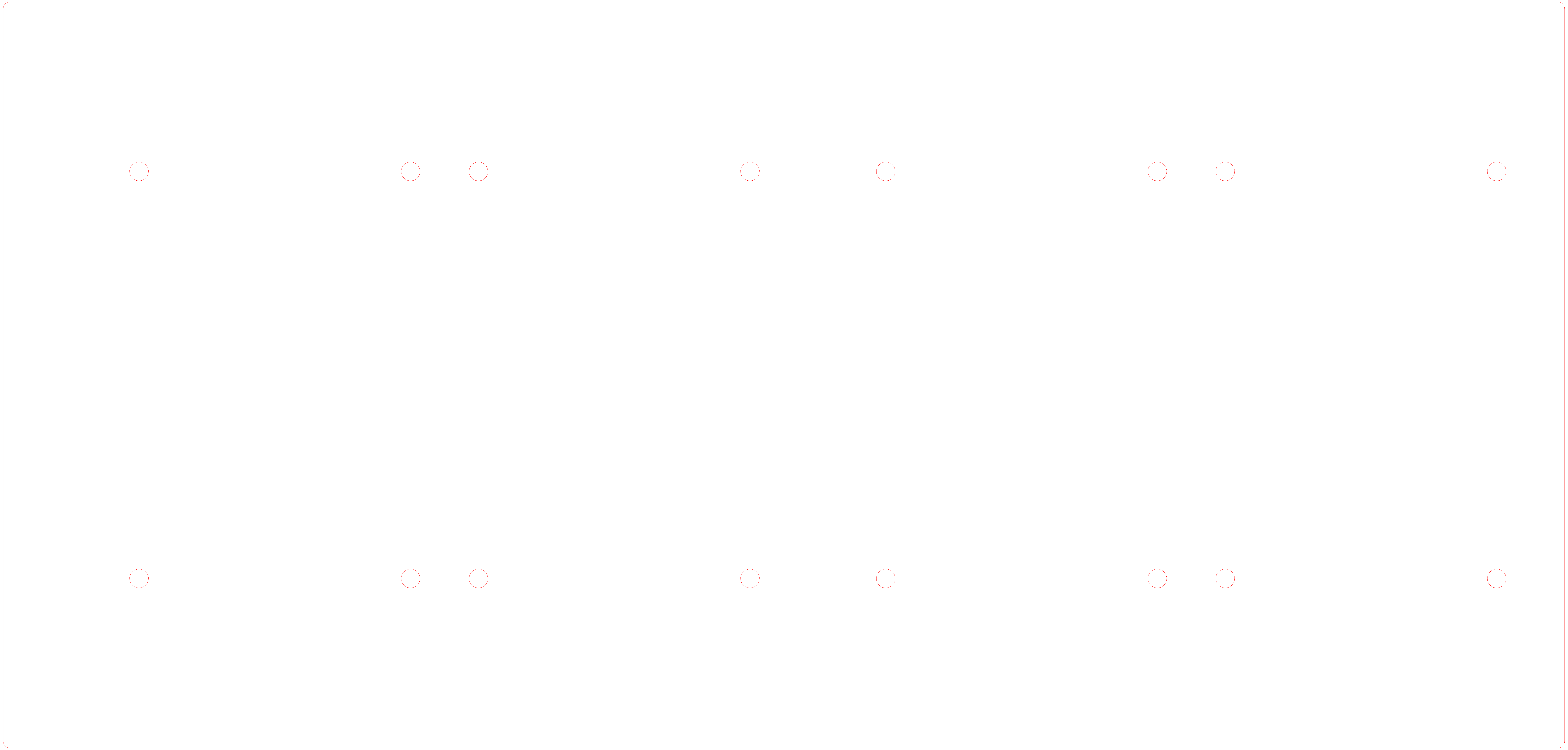

### Basic lifespan machine media tray slots 2.0- cut sheet.PDF

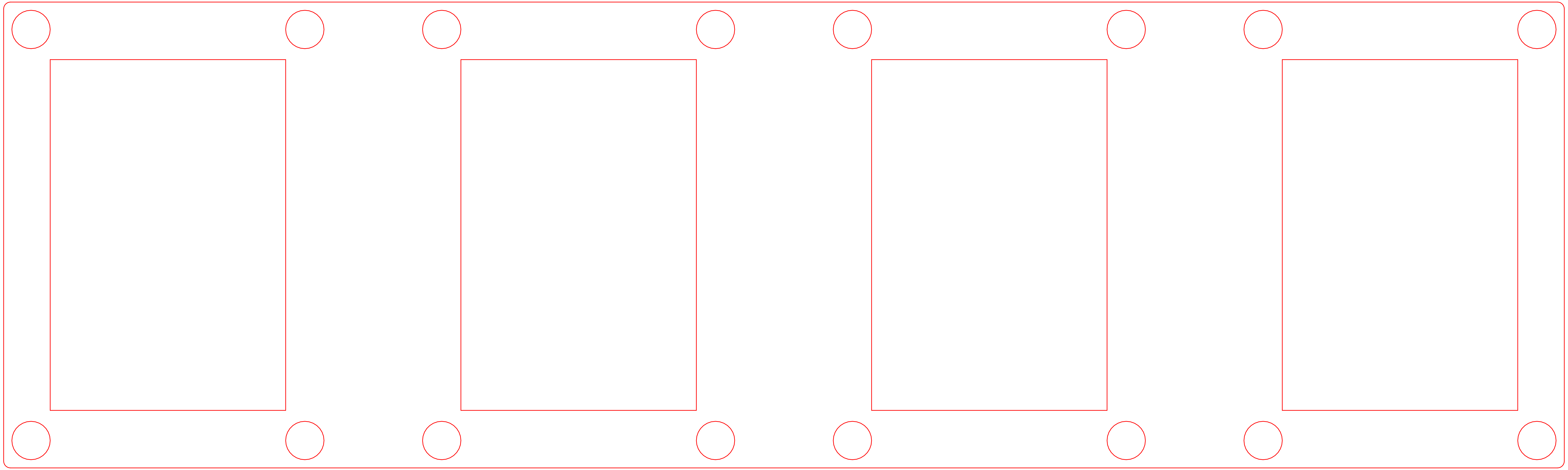

### Basic lifespan machine media tray slots and lip 1.0 - cut sheet.PDF

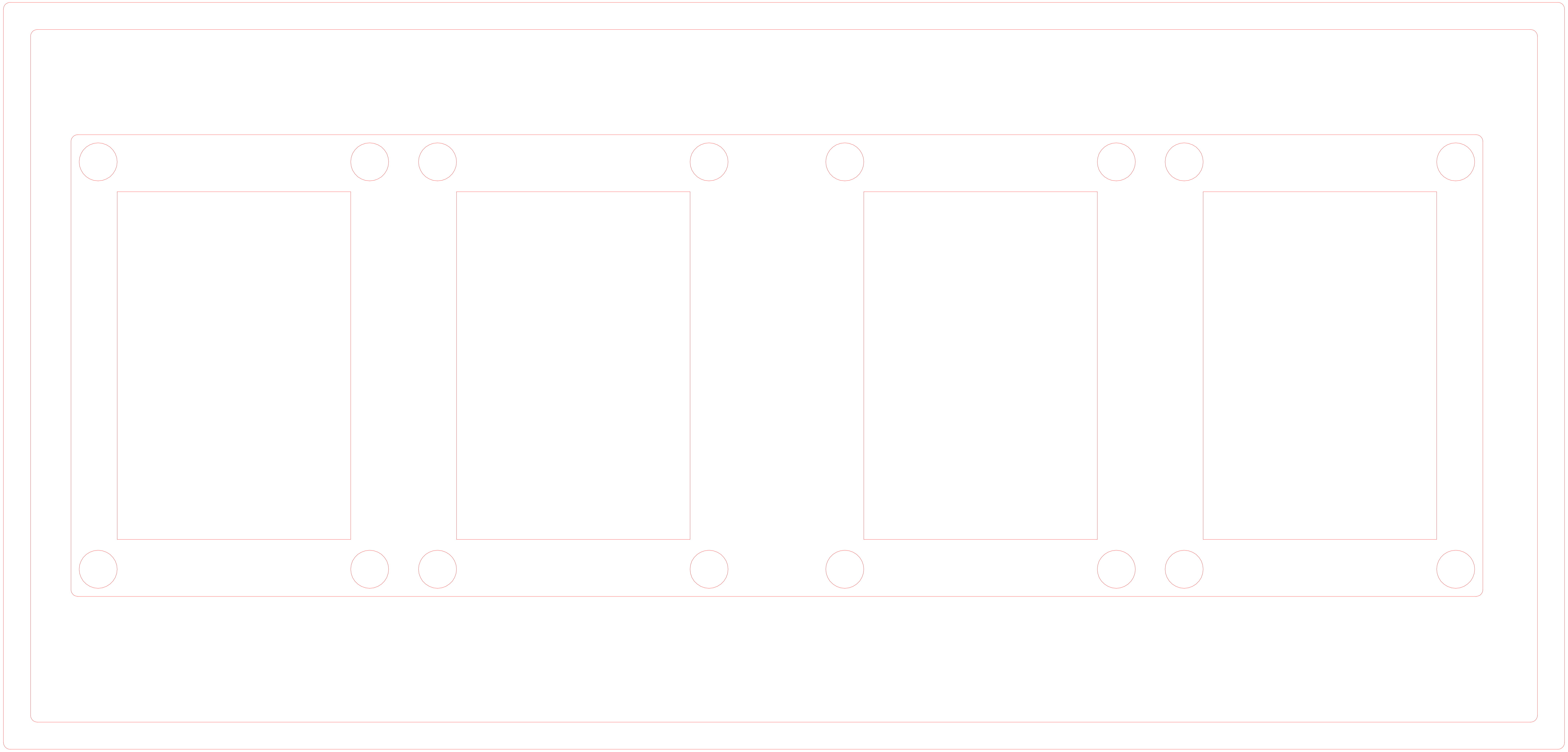

### Supplemental Figure 1

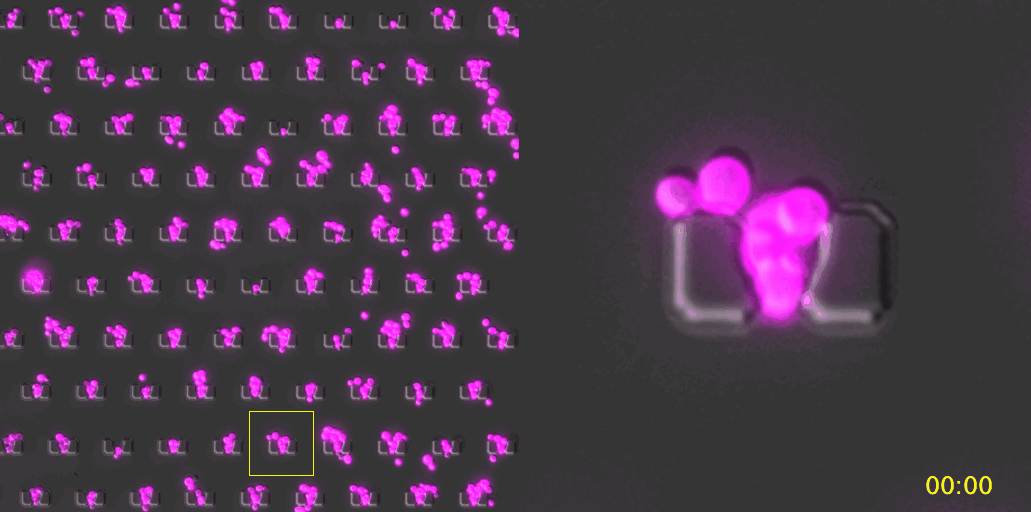
