## Supplementary material for "The Yeast Lifespan Machine: a microfluidic platform for automated replicative lifespan measurements": Designs and Schematics: 24 Well Base Plate.PDF

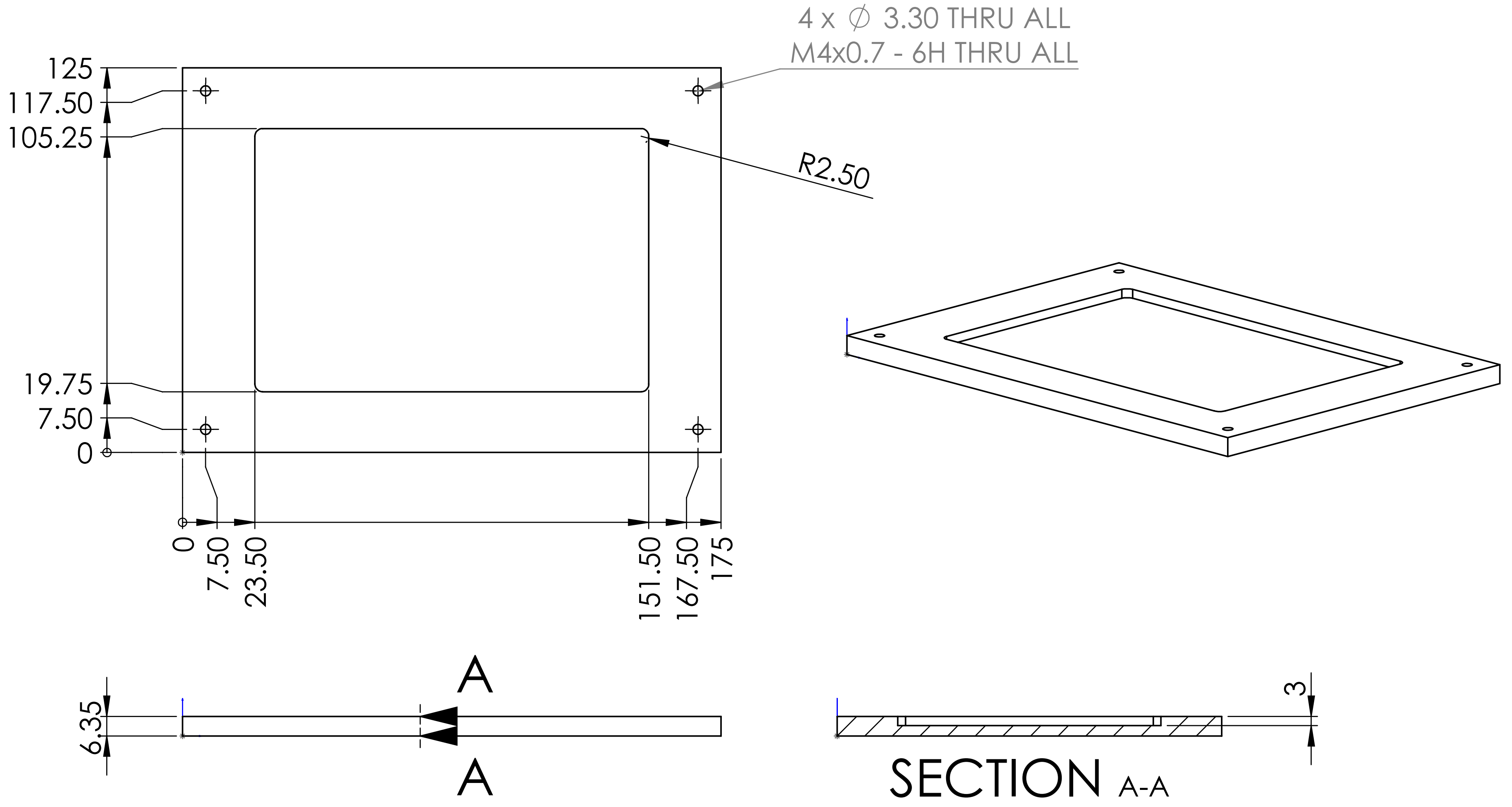


UNLESS OTHERWISE SPECIFIED: DIMENSIONS ARE IN MILLIMETERS

DIMENSIONS ARE IN MILLIMETERS
TOLERANCES:
TWO PLACE DECIMAL ± 0.01

SMOOTH AND DEBUR EDGES

MATERIAL
6061 Aluminum

DO NOT SCALE DRAWING

Calico Life Sciences


CONTACT:
Robert Keyser

650-267-7923


| SIZE | DWG. NO. | REV |
|---|---|---|
| A | 24 WELL BASE PLATE | |

SCALE: 1:2 | SHEET 1 OF 1
