## Supplementary material for "The Yeast Lifespan Machine: a microfluidic platform for automated replicative lifespan measurements": Designs and Schematics: 24 Well Manifold.PDF

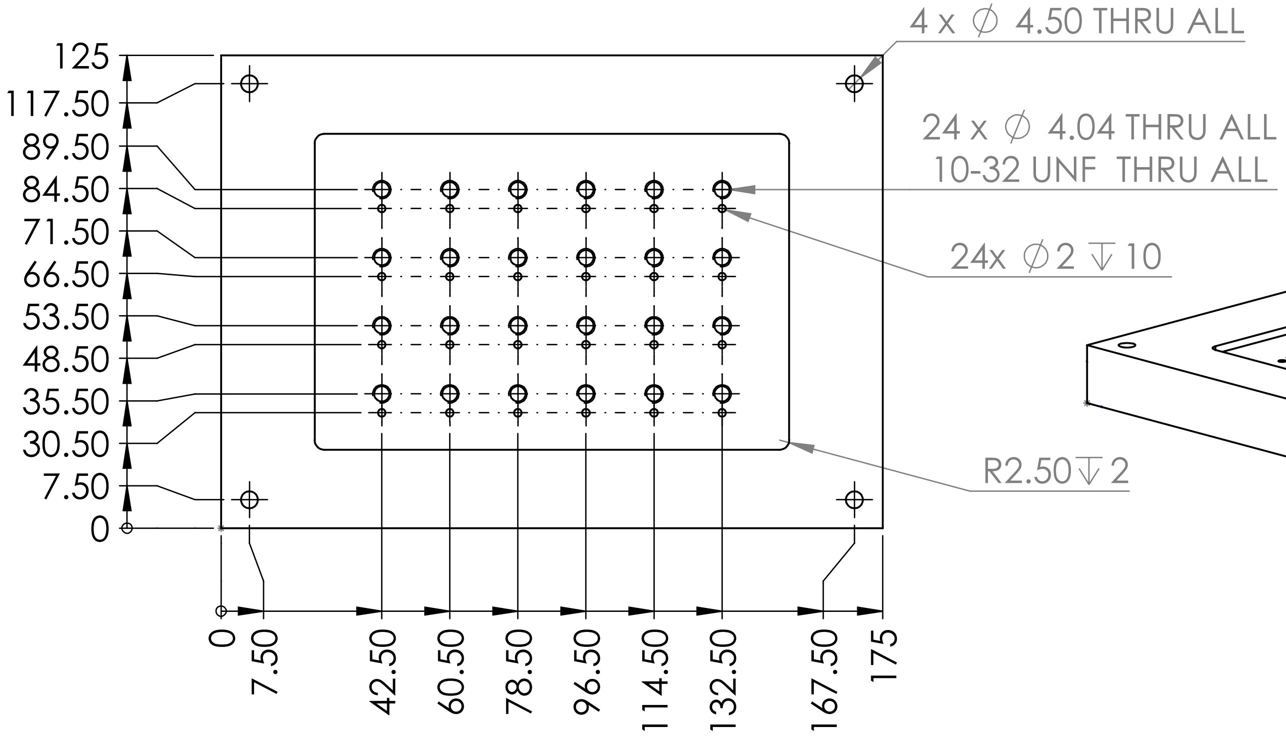


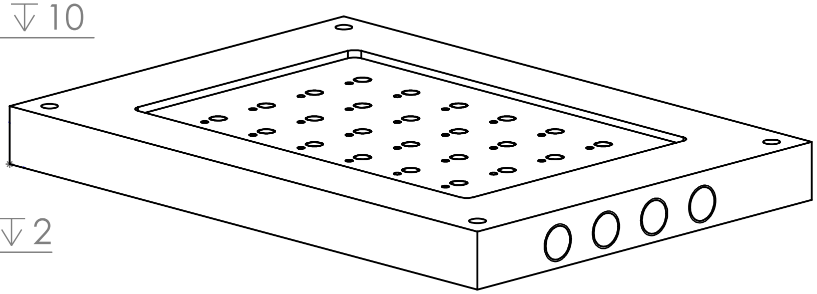

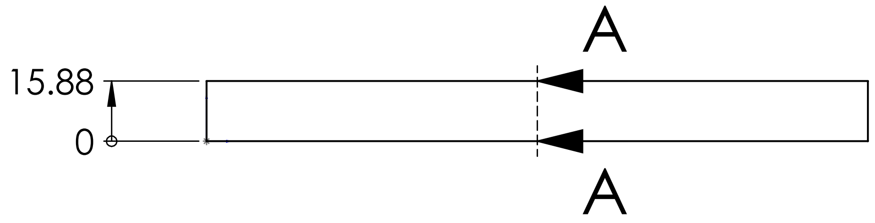


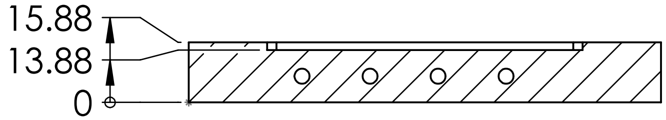


SECTION A-A

UNLESS OTHERWISE SPECIFIED: DIMENSIONS ARE IN MILLIMETERS

DIMENSIONS ARE IN MILLIMETERS
TOLERANCES: ± 0.1mm

SMOOTH AND DEBUR EDGES

MATERIAL
6061 Aluminum

DO NOT SCALE DRAWING

Calico Life Sciences


CONTACT:
Robert Keyser

650-267-7923


SIZE A | DWG. NO. 24WELL VAC MANIFOLD | REV

SCALE: 1:2

SHEET 1 OF 2

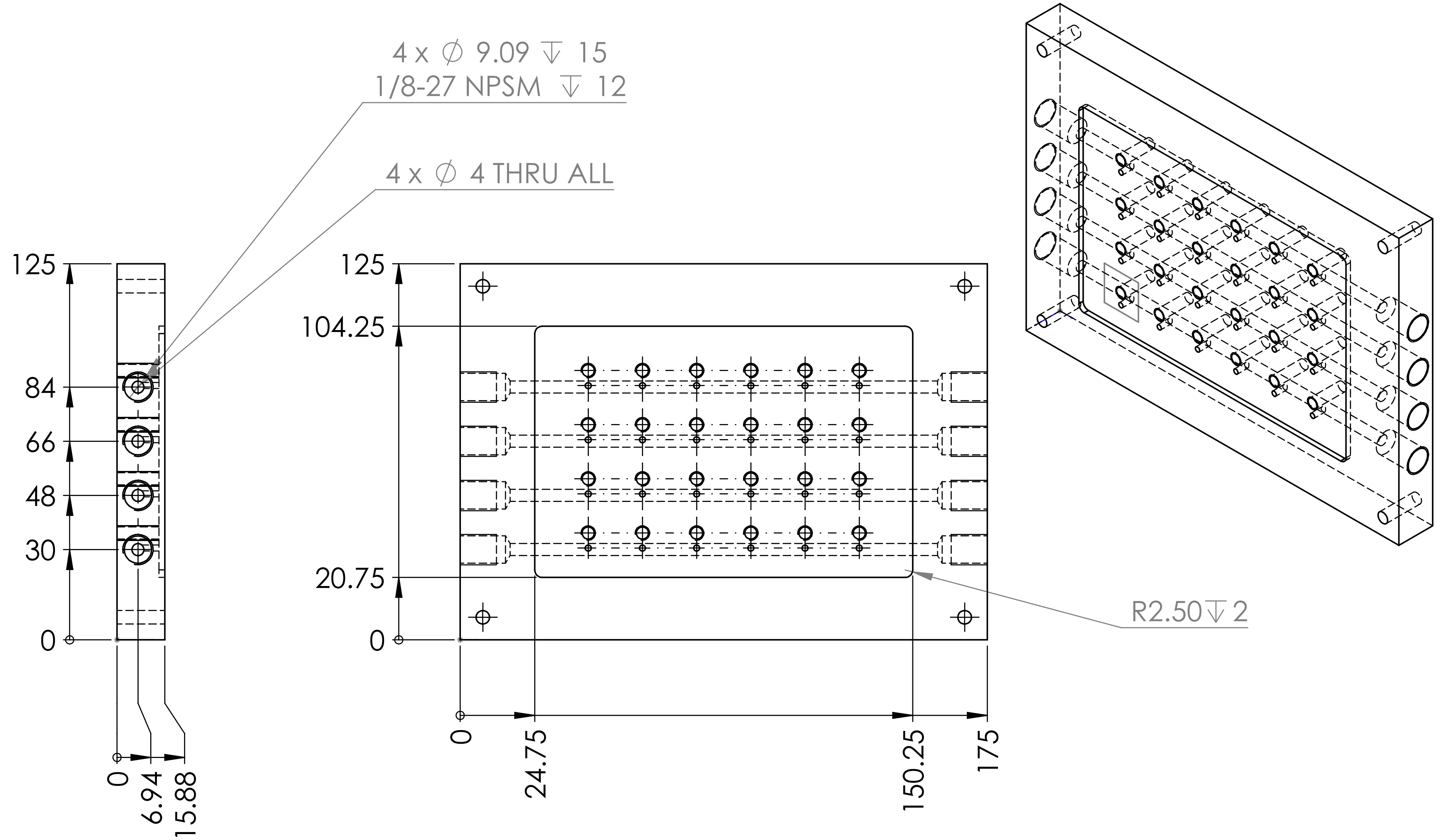


UNLESS OTHERWISE SPECIFIED: DIMENSIONS ARE IN MILLIMETERS

DIMENSIONS ARE IN MILLIMETERS
TOLERANCES: ± 0.1mm

SMOOTH AND DEBUR EDGES

MATERIAL
6061 Aluminum

DO NOT SCALE DRAWING

Calico Life Sciences

CONTACT:
Robert Keyser

650-267-7923

| SIZE | DWG. NO. | REV |
|---|---|---|
| A | 24WELL VAC MANIFOLD | |

SCALE: 1:2
