## Supplementary material for "The Yeast Lifespan Machine: a microfluidic platform for automated replicative lifespan measurements": Designs and Schematics: Device Holder Bottom.PDF

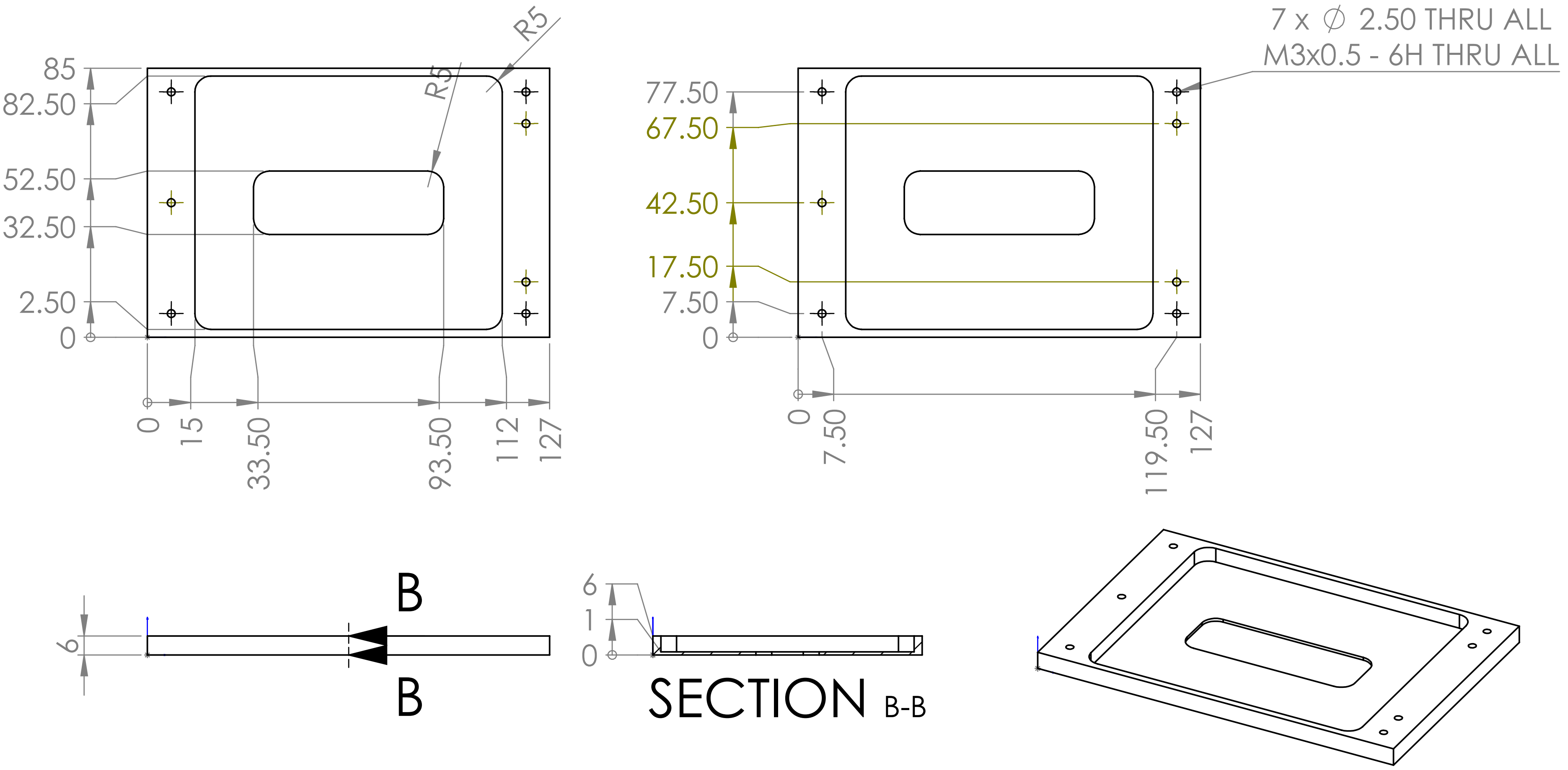


| | |
|---|---|
| UNLESS OTHERWISE SPECIFIED: DIMENSIONS ARE IN INCHES | Calico Life Sciences |
| DIMENSIONS ARE IN MM<br>TOLERANCES: ± 0.01 | CONTACT:<br>Robert Keyser<br><br>650-267-7923 |
| SMOOTH AND DEBUR EDGES | |
| MATERIAL<br>6061 Aluminum | SIZE A · DWG. NO. THIN BOTTOM FRAME · REV |
| DO NOT SCALE DRAWING | SCALE: 1:2 · SHEET 1 OF 1 |
