## Supplementary material for "The Yeast Lifespan Machine: a microfluidic platform for automated replicative lifespan measurements": Designs and Schematics: Media reservoir base plate 6x50ml.PDF

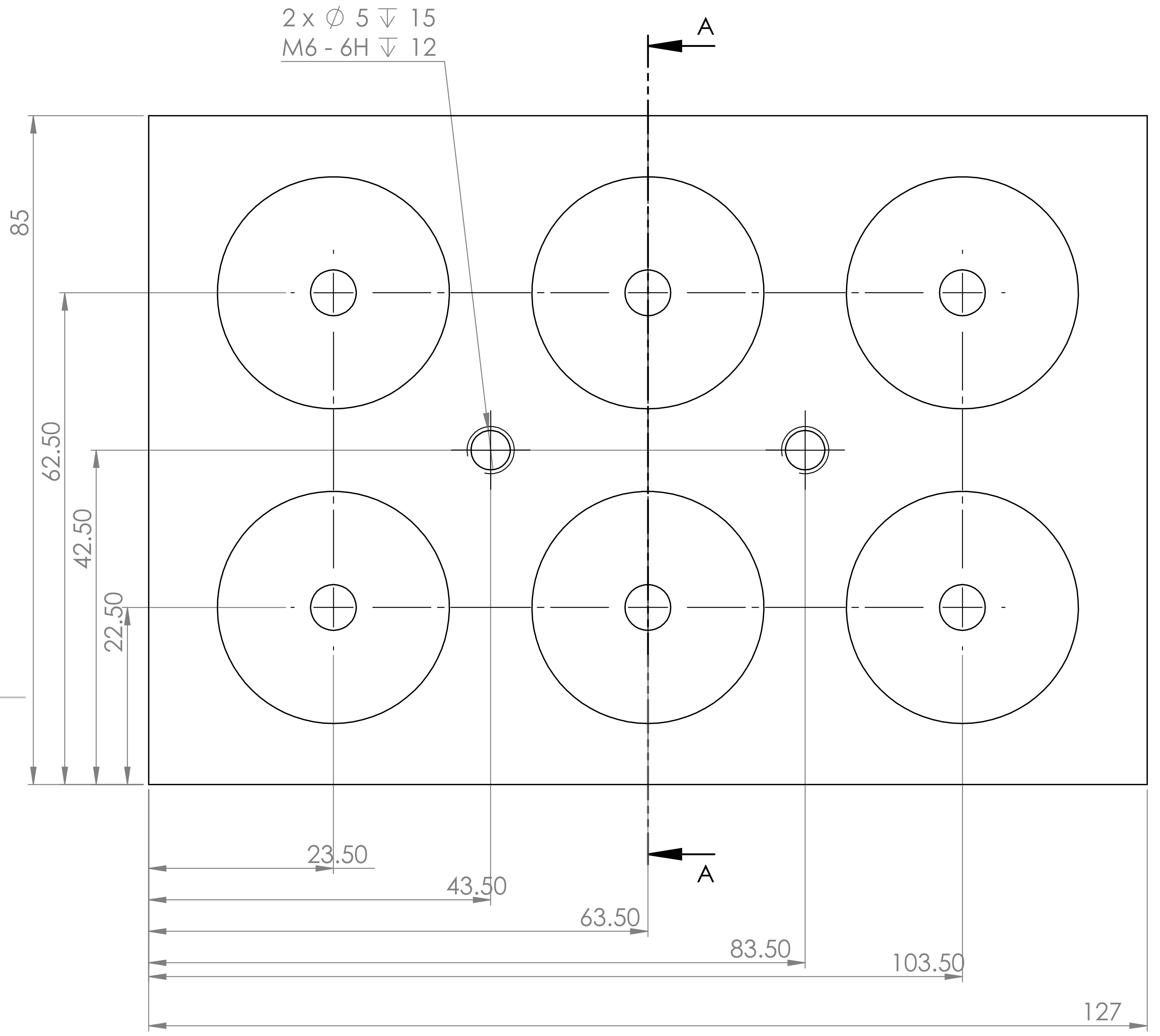


SECTION A-A
SCALE 1.5 : 1

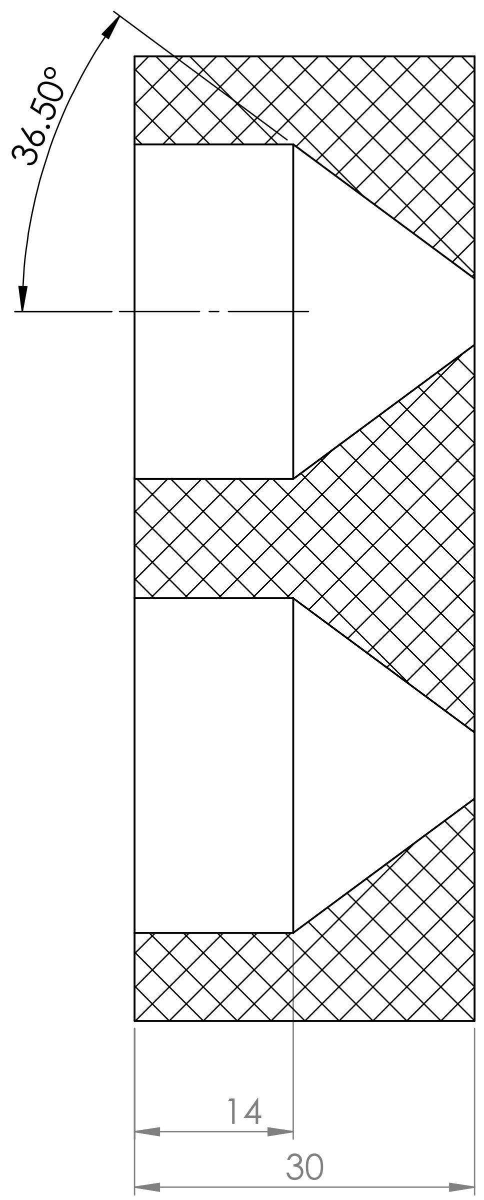


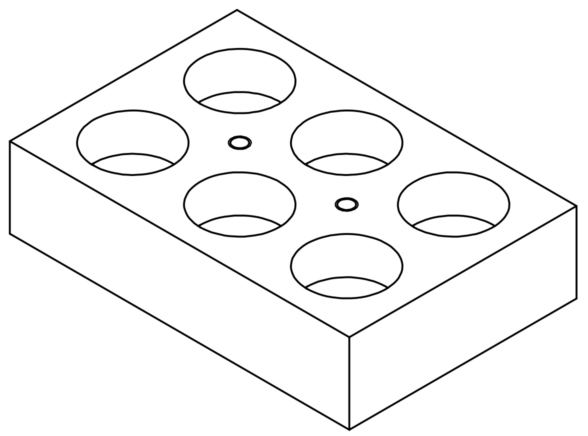

UNLESS OTHERWISE SPECIFIED:
DIMENSIONS ARE IN MILLIMETERS
SURFACE FINISH: N7 = Ra-1.6um-
TOLERANCES:
LINEAR: ±0.1mm
ANGULAR: ±0.5°

FINISH:

DEBURR AND BREAK SHARP EDGES

DO NOT SCALE DRAWING

REVISION A

Calico Life Sciences LLC
1170 Veterans Blvd, South San Francisco, CA, 94080
+1-650-754-6203
www.calicolabs.com

Calico

| | NAME | SIGNATURE | DATE |
|---|---|---|---|
| DRAWN | A.M.S. | | 1/4/2019 |
| CHK'D | | | |
| APPV'D | | | |
| MFG | | | |
| Q.A | | | |

Description:

MATERIAL:

File name:
Media reservoir holder 6x50ml base plate 1.2

A3

WEIGHT:

SCALE:1:1

SHEET 1 OF 1
