## Supplementary material for "The Yeast Lifespan Machine: a microfluidic platform for automated replicative lifespan measurements": Designs and Schematics: Media reservoir manifold 6x50ml.PDF

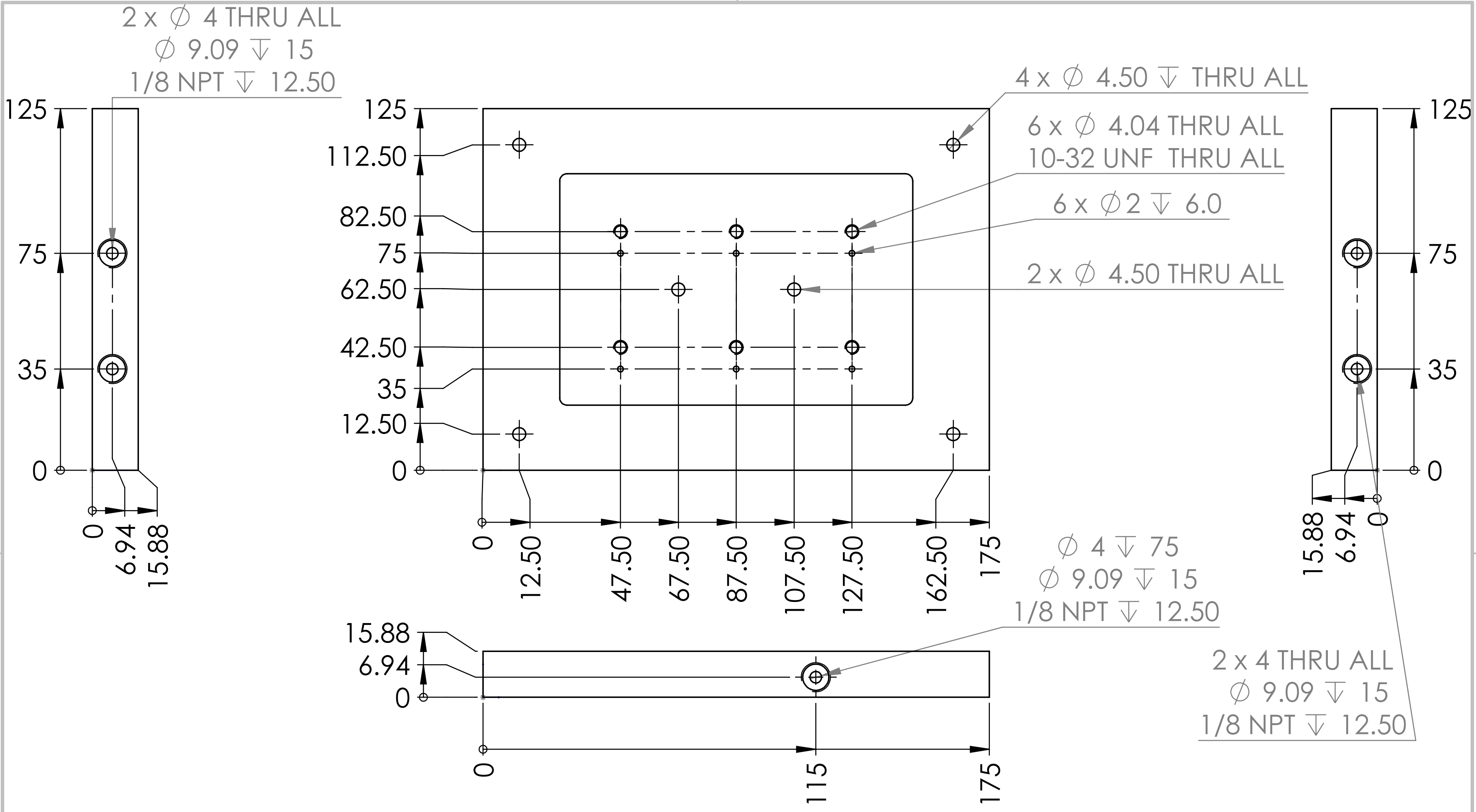


| | |
|---|---|
| UNLESS OTHERWISE SPECIFIED: DIMENSIONS ARE IN MILLIMETERS | Calico Life Sciences |
| DIMENSIONS ARE IN MILLIMETERS<br>TOLERANCES: ± 0.1 | CONTACT:<br>Robert Keyser<br><br>650-267-7923 |
| SMOOTH AND DEBUR EDGES | |
| MATERIAL<br>6061 Aluminum | SIZE A — DWG. NO. 6X50ML PLATE 004 — REV |
| DO NOT SCALE DRAWING | SCALE: 1:2 — SHEET 1 OF 2 |

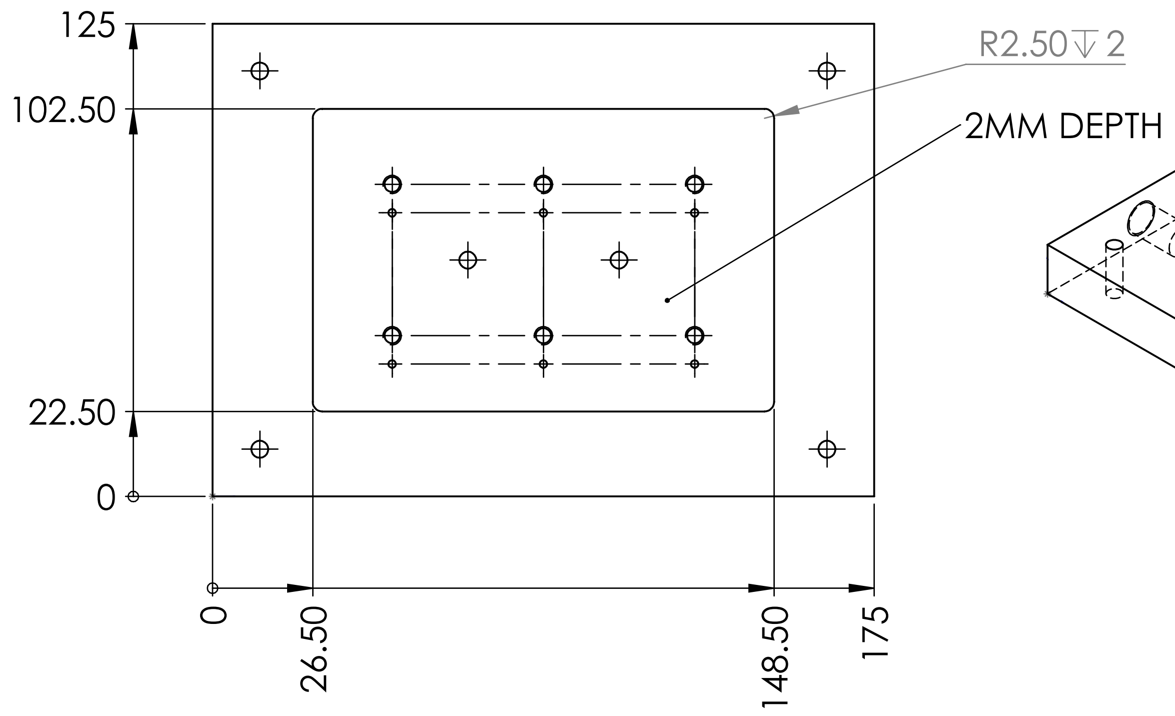


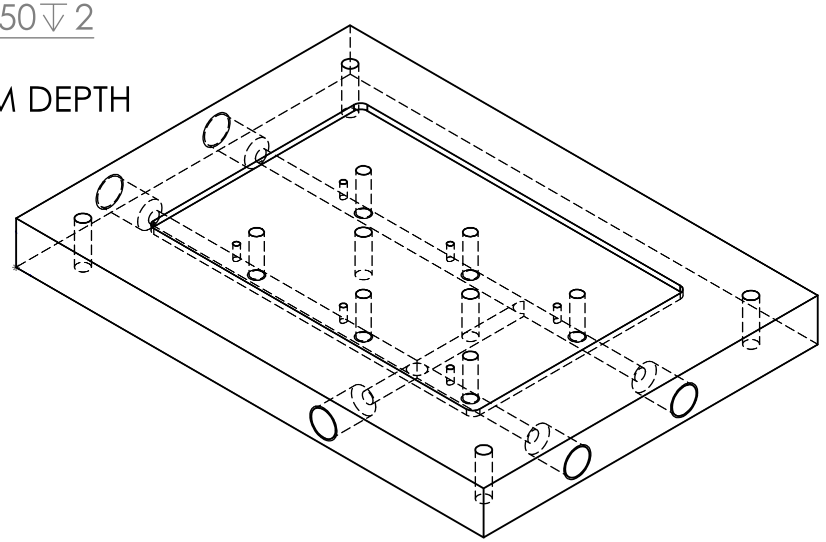

| | |
|---|---|
| UNLESS OTHERWISE SPECIFIED: DIMENSIONS ARE IN MILLIMETERTS | Calico Life Sciences |
| DIMENSIONS ARE IN MILLIMETERS<br>TOLERANCES: ± 0.1 | CONTACT:<br>Robert Keyser<br><br>650-267-7923 |
| SMOOTH AND DEBUR EDGES | |
| MATERIAL<br>6061 Aluminum | SIZE A | DWG. NO. 6X50ML PLATE 004 | REV |
| DO NOT SCALE DRAWING | SCALE: 1:2 | SHEET 2 OF 2 |
